# Big data analysis of animal movements in aquatic ecosystems with acoustic telemetry

**DOI:** 10.64898/2026.06.01.729394

**Authors:** Edward Lavender, Matthew H. Futia, Andreas Scheidegger, Stanisław W. Biber, Jakob Brodersen, Robert A. Briers, James Thorburn, Carlo Albert

**Affiliations:** School of Applied Sciences, Edinburgh Napier University, UK; Centre for Conservation and Restoration Science, Edinburgh Napier University, UK; Department of Systems Analysis, Integrated Assessment and Modelling, Eawag Swiss Federal Institute for Aquatic Science and Technology, Switzerland; Rubenstein Ecosystem Science Laboratory, University of Vermont, US; School of Mathematics and Physics, University of Surrey, UK; Department of Fish Ecology and Evolution, Eawag Swiss Federal Institute for Aquatic Science and Technology, Switzerland

**Keywords:** Bayesian inference, biotelemetry, geolocation, management, movement ecology, particle filter, *Salvelinus namaycush*, state-space modelling

## Abstract

Underwater receiver networks (passive acoustic telemetry systems) are deployed to track animals in aquatic habitats all over the world, but remarkably limited attention has been given over to how we can strengthen the value of these networks through statistical and computational advances. Here, we upscale state-of-the-art methods of Bayesian inference to big animal-tracking datasets from acoustic telemetry, with the largest geolocation analysis in a sparse passive acoustic telemetry system (with non-overlapping receivers) to date. Using four years of data from 93 lake trout (*Salvelinus namaycush*) in North America’s Lake Champlain (657,360 timesteps per individual), we formulate and fit state-space models to reconstruct animal movements through time. Uniquely, we directly embed biological expertise and detailed complementary datasets from fine-scale positioning systems, accelerometry, swim-tunnel experiments and field range tests in our analysis. Using simulated and real-world datasets, we map movement patterns and estimate residency in distinct management zones. We quantify array precision and deliver maps and residency estimates with a median error and precision (standard error) below 1 %. These results strengthen the evidence base for management. This work takes us a step towards robust inference of movement patterns at scale in acoustic telemetry systems across the world. We can, and should, build on prior scientific progress and extend the value of hard-earned data beyond individual studies to refine inferences for ecology and management.

**Significance statement:** Acoustic receivers are deployed across the globe to track aquatic animals, but reconstructing detailed movement patterns from detections at receivers remains a considerable challenge. Here, we upscale state-of-the-art methods of Bayesian inference by two orders of magnitude to analyse big, real-world datasets, using an extensive case study of lake trout (*Salvelinus namaycush*) in Lake Champlain. By directly integrating diverse complementary datasets from animal-borne tags, swim-tunnel experiments, field studies and close-kin mark-recapture in our analysis, we resolve detailed movement patterns over a four-year period, with broad implications for ecology and management. This work provides a powerful framework for acoustic telemetry studies that strives to meet the challenges of big, real-world datasets from telemetry networks across the world.

## 1. Introduction

We are in the midst of a revolution in animal tracking (1, 2). Electronic tagging and tracking technologies, especially satellite-based trackers, are being deployed to track or geolocate animals across the globe, providing unprecedented insights into animal movements (3), space use (4) and behaviour (5), with wide-ranging implications for biology (6), environmental science (7) and wildlife conservation (8). However, animal geolocation in habitats that satellite signals cannot penetrate, especially underwater, remains challenging (9).

In aquatic habitats, passive acoustic telemetry systems are extensively deployed (10). These systems comprise networks of receivers (hydrophones), which listen for acoustic transmissions from tagged animals. The transmitters are programmed to release regular, high-frequency, individual-specific pings, which can be recorded by receivers when animals swim within range (11). In fine-scale arrays where receiver ranges are overlapping (hereafter ‘acoustic positioning systems’), simultaneous detections of tagged animals at multiple receivers enable high-resolution tracking. However, in many (hereafter ‘sparse’) passive acoustic telemetry systems receiver ranges are non-overlapping. These systems provide important data on animal presence around receivers (12), but reconstructing individual movements from sparse detections has remained a significant hurdle, hindering ecological analyses (13) and management applications (14). To date, most studies have relied on heuristic approaches (15), which generally smooth over detection (receiver) locations or interpolated ‘pseudo-positions’ to generate maps of space use without uncertainty quantification (16). These approaches do not formally geolocate animals and can indicate erroneous maps of space use (17, 18).

From a statistical perspective, the challenge is to derive the posterior distribution *f*(***s***_1:*T*_ | ***y***_1:*T*_) for the unknown trajectories ***s***_1:*T*_ of a tagged animal, given the observations ***y***_1:*T*_ (16). We use the notation *f*(·) to denote a probability density function, ***s***_*t*_ to denote the unknown state (which includes the two-dimensional location) of an animal at time *t*, ***y***_*t*_ to denote acoustic observations (detections, non-detections) at receivers and *t* = 1, 2, …, *T* to index time. In an acoustic telemetry system, two processes provide information: a movement process describing our prior knowledge of the animal’s movement behaviour and an observation (detection) process that connects the animal’s position to the observations. Using Bayes’ Theorem, we can express the posterior distribution in terms of these processes as *f*(***s***_1:*T*_ | ***y***_1:*T*_) ∝ *f*(***s***_1:*T*_) *f*(***y***_1:*T*_ | ***s***_1:*T*_), where *f*(***s***_1:*T*_) is a model of the movement process and *f*(***y***_1:*T*_ | ***s***_1:*T*_) is a model of the observation process. Together, the two models form a Bayesian state-space model (19).

By performing inference for the state-space model, we can approximate *f*(***s***_1:*T*_ | ***y***_1:*T*_). That is, we can estimate an animal’s unknown locations and uncertainty, while leveraging our biological knowledge about *f*(***s***_1:*T*_) and *f*(***y***_1:*T*_ | ***s***_1:*T*_) in the analysis. For example, accelerometery datasets (20), fine-scale positioning analyses (21) and domain expertise can be embedded in the movement model. Similarly, we can use studies of receiver detection ranges (22) to derive data-driven acoustic observation models, while accounting for physical considerations and hardware specifications. These studies are widely conducted but almost never integrated in acoustic telemetry analyses, despite extensive recognition of their importance (11).

The development of state-space models for animal geolocation that integrate biological knowledge is critical to advance the use of acoustic telemetry in ecology and management (16). Robust, probabilistic estimates of individual locations are the foundation of downstream ecological analyses and vital for our understanding of the ecology and management requirements of tracked species (16). However, few studies have adopted this approach (18, 23–26). Furthermore, existing studies have focused almost exclusively on illustrative simulation (18, 25) or example analyses (23, 24), with few individuals and short datasets. There is a great gulf between the scale of these analyses and real-world datasets (27).

Here, we present the largest geolocation analysis in a sparse passive acoustic telemetry system to date. Focusing on a case study of lake trout (*Salvelinus namaycush*) in North America’s Lake Champlain (28), we analyse a dataset comprising 1,735,137 detections of 93 individuals, derived from 153 receiver deployments in 31 stations over a four-year (657,360 timestep) period (Fig. 1). Uniquely, we directly build on previous research, integrating extensive prior knowledge and data in our analysis (Fig. 2). This approach embodies the nature of scientific progress (29), demonstrating how we can analyse big, real-world datasets with probabilistic models and in so doing strengthen the value of animal-tracking data for ecology and management. This work advances our understanding of animal movements and provides a quantitative basis for management in acoustic telemetry systems across the world (30).

**Fig 1.**
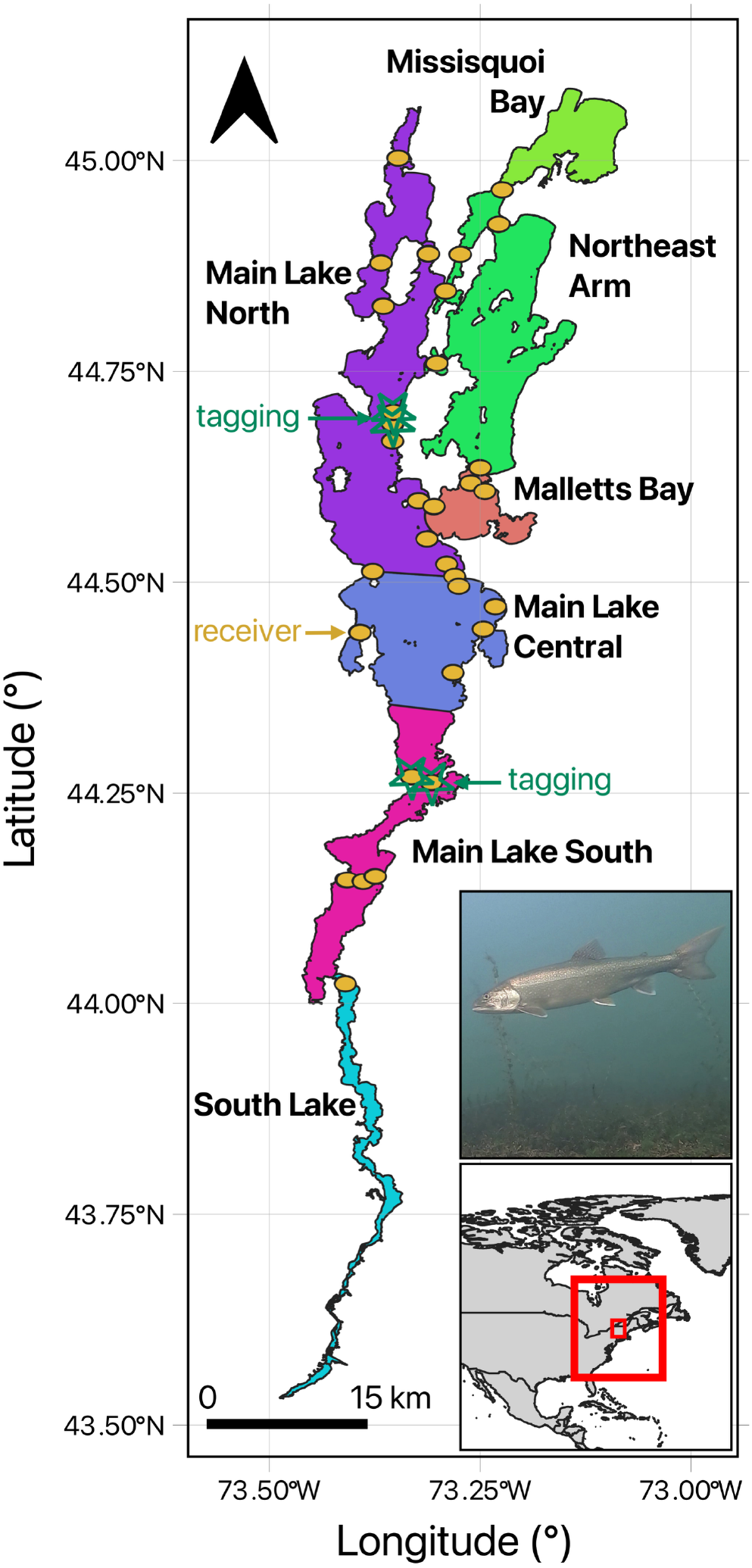
The study system. The insets show the location of Lake Champlain in North America and a lake trout from this system. The main panel shows the lake, split into seven regions following ref. (28), lake trout tagging locations and receivers (point diameter: 1000 m). Spatial data sourced from refs. (31, 32). Photo credit: Matthew Futia.

**Fig 2.**
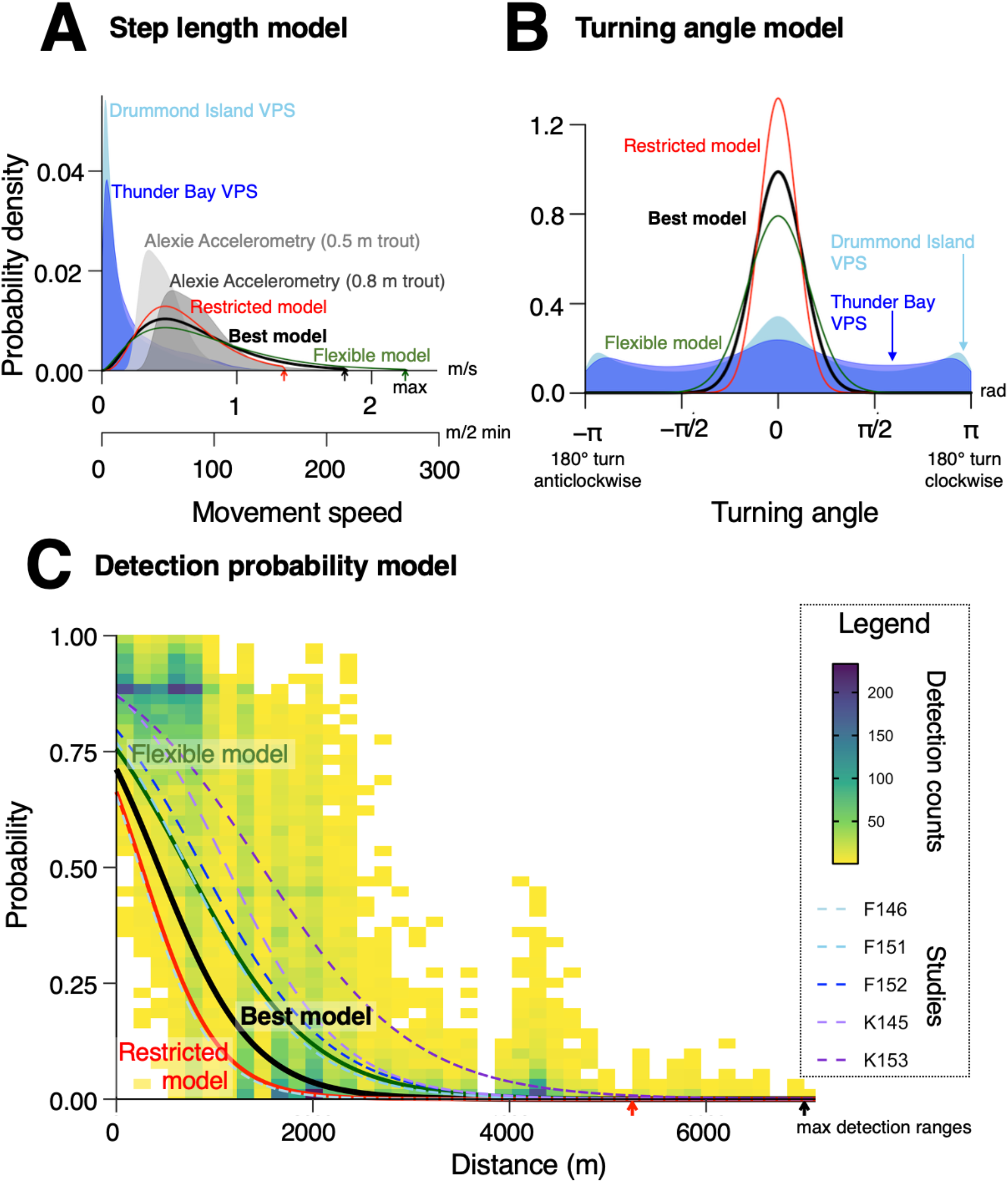
A state-space model for lake trout in Lake Champlain. This comprises a movement model (**A–B**) and an observation model (**C**). All models were informed by literature, complementary datasets, domain knowledge and a model validation exercise (see Methods and Supporting Information §1–4). The temporal resolution was two minutes. We derived a best parameterisation (black) plus restrictive (red) and flexible (green) parameterisations (for sensitivity analyses). **A** shows truncated Gamma step length models alongside empirical distributions derived from three datasets, labelled Drummond Island VPS (21), Thunder Bay VPS (33) and Alexie Accelerometry (20, 34). Arrows mark the maximum movement speed (mobility). **B** shows mixture (normal-uniform) turning angle models alongside distributions from positional telemetry (21, 33). **C** shows truncated logistic distance-decay models for detection probability at acoustic receivers, conditional on a line of sight between receiver and transmitter. Detection probability was set to zero if the line of sight was blocked by land. Arrows mark the truncation parameter (*γ*). Models were informed by range-testing datasets from Futia et al. (35) and Klinard et al. (22). Dashed lines show empirical detection probability relationships for each study (F, K) and transmitter power (145–153 dB).

## 2. Results

### 2.1 Study system

We analysed lake trout movement patterns in Lake Champlain (Fig. 1, Tables S1–2). To do so, we formulated a state-space model and then performed locational inference using Bayesian particle filtering–smoothing algorithms (Fig. 2, Tables S3–4). For our workflow, see Fig. S1. For the lake trout dataset, see Fig. S2.

### 2.2 Simulation analysis

To begin, we reconstructed movements for a synthetic dataset of 96 simulated individuals at two-minute resolution over a one-month period to examine algorithm performance and sensitivity. The number of days with detections ranged from 2–31 (median = 26). Our main analysis used the same model formulation for simulation and inference. In this analysis, we successfully reconstructed movements for all individuals. There was a close correspondence between simulated trajectories and inferred occurrence distributions (Figs 3 and S3). The median spatial uncertainty in individual locations (the median area spanned by 95 % of the probability mass) was 124 cells (0.4 % of the lake area). Residency within each of the lake regions was estimated within an accuracy of ± 6 %, which exceeds the accuracy of a heuristic (interpolation) approach we included for comparison (Figs 3 and S4). Across all individuals, the Mean Weighted Occupancy Error (MOE) over all regions ranged from 0.0–5.2 (median = 0.7) %, which compares to 0.0–99.3 (median = 4.9) % for the heuristic approach. The standard deviation (error) in MOE (a measure of array precision) was 0.99 % versus 18.6 % for the heuristic approach (Fig. S5).

**Fig 3.**
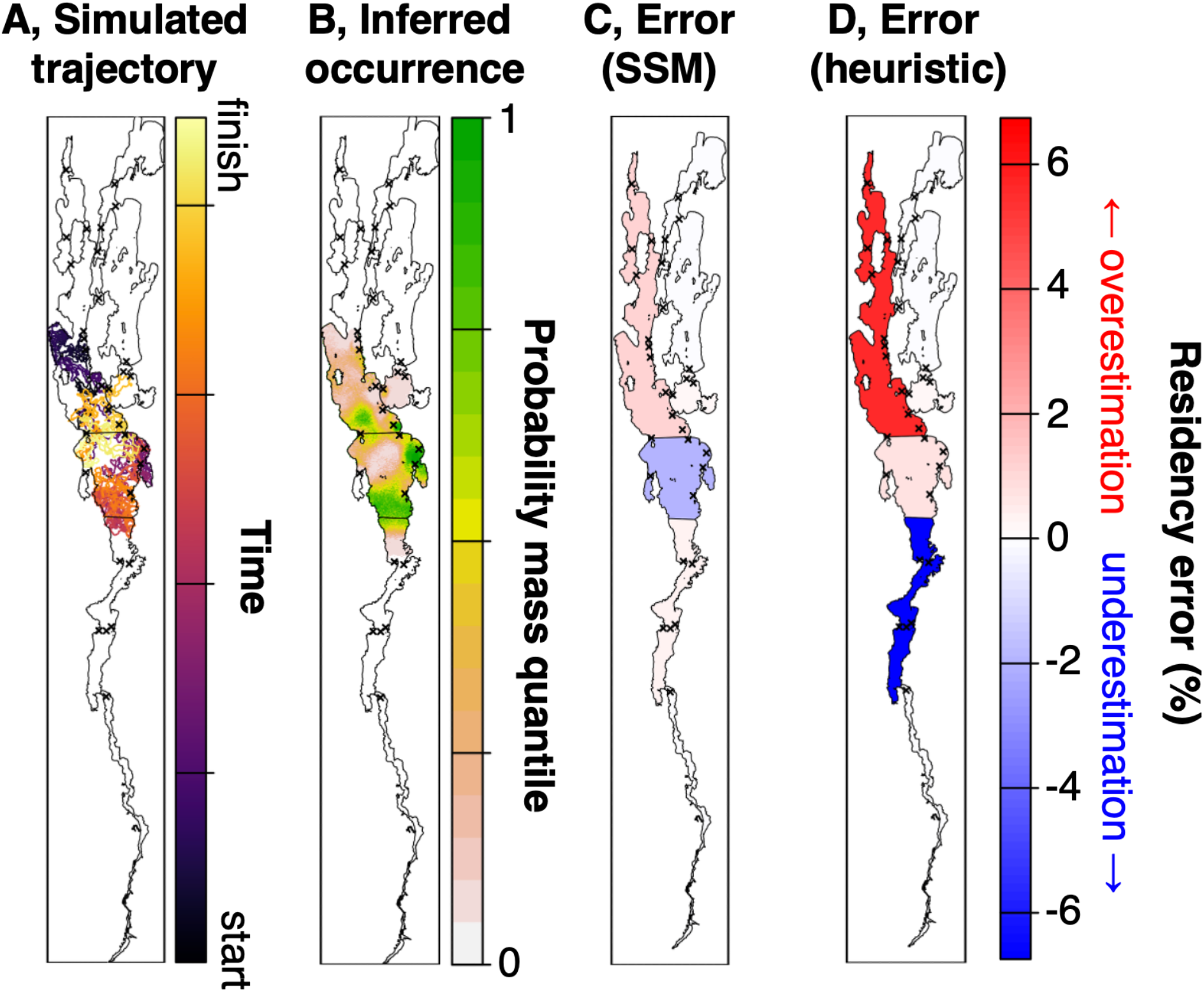
Simulation analyses. **A** shows a simulated trajectory. **B** shows the inferred occurrence distribution. Colours show probability quantiles; the 0.5 and 0.95 contours enclose the regions containing 50 % and 95 % of the probability mass. **C** shows the difference between the simulated and estimated percentage of time steps spent in each region from state-space modelling. **D** shows the same metric for a heuristic (interpolation) method included for comparison. Crosses mark receiver locations.

To analyse sensitivity, we re-ran the inference process with restrictive and flexible parameterisations of the movement (step length, turning angle) and observation (detection probability) models (96 × 2 × 3 = 576 algorithm runs). In this analysis, 99 % of runs converged. Convergence failures were driven by misspecification of the acoustic observation model, with the overly flexible detection probability function effectively blocking movements that occurred without detection through receiver barriers. For successful runs, occurrence distributions and residency estimates were robust to the degree of parameter misspecification we explored (Figs 3 and S3–5).

Across all 672 runs, total single-threaded computation time ranged from 137–179 (median = 149) minutes per dataset (Fig. S6). This includes filtering (11 minutes) with 10,000 particles (× 2) and two-filter smoothing (126 minutes) with 1,500 particles. The median effective sample size after smoothing was 123 particles (Fig. S6). Proper smoothing was achieved on 93–100 % of time steps.

#### 2.3. Real-world analysis

In the real-world analysis, the number of days with detections ranged from 1–31 (median = 10) per month. Lake trout distributions varied by tagging location and season (Figs 4 and S7). In fall (October–November), there were clear differences between the two tagging groups, with north-tagged fish more resident in North Main Lake (plus Central Main Lake and Northeast Arm) and south-tagged fish more resident in South Main Lake. Thereafter, the two groups exhibited similar seasonal trends. During winter (December–March) and spring (April–May), both groups exhibited high residency in Main Lake North, Malletts Bay, Main Lake Central and South Main Lake. During winter, south-tagged fish also exhibited elevated residency in South Lake. Throughout fall to spring, there were some apparent hotspots in bays without receivers. Over summer (June–September), movements concentrated in South and Central Main Lake, restricted by the thermal habitat suitability threshold in the model. North-tagged fish also showed relatively higher residency in North Main Lake and Northeast Arm during this time.

**Fig 4.**
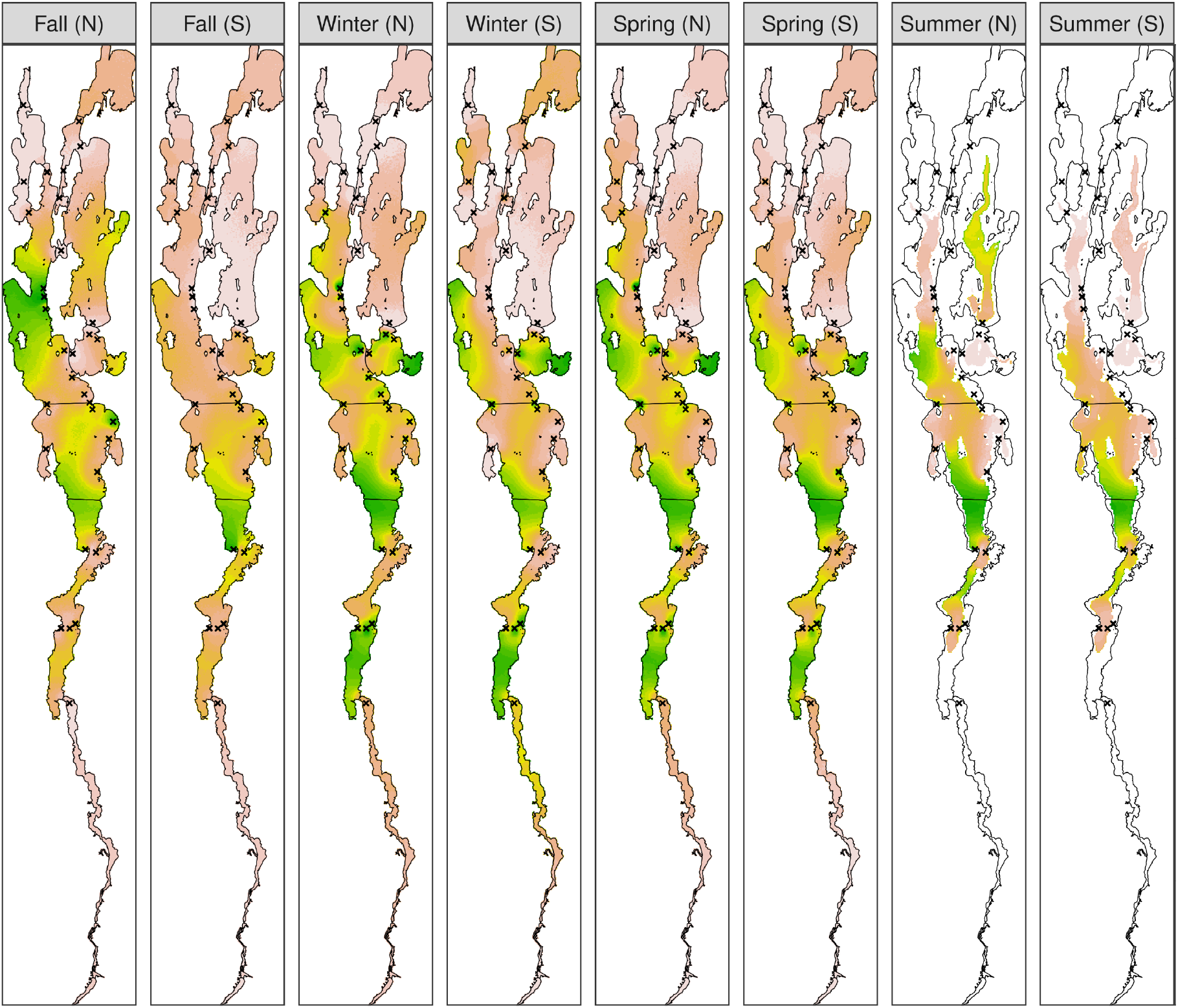
Lake trout space use in Lake Champlain. Each map is the overall probability distribution for the location of modelled lake trout by tagging site (North, N or South, S) and season. Map properties follow Fig. 3.

The analysis was implemented in overlapping blocks of approximately monthly duration (20,160–58,572 time steps). Convergence rates for the particle filter and smoother were 78 % and 91 % (after successful filtering) respectively. Convergence was poorest for summertime blocks (52 %), when detection time series were poorest, and in the south of the lake where individuals sometimes passed through receiver barriers without detection. For successful blocks, computation time averaged 535 minutes, including filtering (87 minutes) with 50,000– 100,000 particles (× 2) and two-filter smoothing (425 minutes) with 2,500 particles. After smoothing, the median spatial uncertainty was 266 cells (8 % of the lake area). The median effective sample size was 256 particles (Fig. S8). Proper smoothing was achieved on 75–100 % of time steps.

## 3. Discussion

We have presented the largest geolocation analysis of animal movement patterns in a sparse passive acoustic telemetry system. This work extends previous studies relying on heuristic approaches (15) with a state-space modelling framework (16) that directly integrates prior knowledge, additional datasets and biological expertise. Importantly, this approach is applicable to systems with limited receiver coverage, low (> 60 s) transmission rates and irregular detections. As shown by our lake trout analysis, we can now resolve detailed, system-wide movement patterns, both within and between periods of detection, while accounting for individual movements (36) and detectability (11) and benefiting from complementary datasets (37). The results provide a solid foundation for future ecological analyses (16). By upscaling Bayesian inference to big, real-world datasets, we hope to enable the adoption of this approach for movement ecology in ecosystems across the world.

This work extends previous studies based on heuristic methods, such as interpolated paths (15). These methods typically use summary statistics and tuning parameters to interpolate movement patterns without uncertainty quantification (16). While such methods are useful data-exploration tools (28), they generally do not model the movement or detection processes that generate observations, despite extensive recognition of their importance (17, 25), and can perform poorly. The development of state-space modelling frameworks that represent these processes, allowing us to incorporate biological knowledge and complementary datasets in analyses, is a step forward. These frameworks provide a robust, probabilistic foundation for analyses of movements, space use and residency that should support research in many systems (16).

This study upscales previous state-space modelling efforts to big, real-world datasets. To date, the largest state-space analysis in a sparse passive acoustic telemetry system was a study of flapper skate (*Dipturus intermedius*) in Scotland (14), though larger datasets have been analysed with heuristic approaches and in acoustic positioning systems (38). The skate study analysed 11 individuals at two-minute resolution over a 14-month period (48 individual/month datasets or approximately one million positions). The authors leveraged prior knowledge and literature for model development (39–41), but were limited by the absence of complementary datasets. Other state-space modelling studies have generally focused on method development, with simulation (18, 25, 42) or small-scale (23, 24) analyses of a handful of individuals over hours or days (or longer periods at coarser temporal resolution (26)). Our study of more than 2,000 individual/month time series expands these efforts by two orders of magnitude. We overcame the computational barrier by exploiting an efficient implementation of particle algorithms for inference (43). We focused inference on location and leveraged complementary datasets (20–22, 33–35) plus a validation analysis to inform and fine-tune our model for efficient inference. We generally achieved computation times of 2–10 hours per individual/block (up to 58,572 location estimates), which is substantial but manageable. A cumulatively expensive feature of our model was the incorporation of (horizontal) line-of-sight in the detection probability function, which is unique to this study. Faster computation times are achievable (18, 43) and we hope readers will develop our workflow for efficient analyses in other systems.

The extensive incorporation of complementary datasets is another distinct feature of this study. There is substantial interest in the use of complementary datasets to bolster acoustic telemetry (37) but this is the first acoustic study to leverage such information extensively within a model-based geolocation framework. Importantly, our framework can benefit from diverse data sources, including accelerometry (20), fine-scale positioning (21, 33), swim-tunnel experiments (34) and in-situ/ex-situ range-testing datasets (22, 35). The value of data integration is increasingly recognised in ecology (44) and, in animal-tracking contexts, the use of ancillary datasets to inform priors and/or as part of the likelihood can support the development of refined models, efficient inference and reduced uncertainty (16). By aligning interest in complementary datasets (37) with inference approaches that can explicitly incorporate such information into analyses (16), we aspire to support cumulative scientific progress in this field.

While previous work has illustrated the benefits of model-based approaches (17, 18, 25), simulations remain an important tool for understanding method performance and sensitivity. For Lake Champlain, our simulations show that we can reconstruct occurrence distributions with notable accuracy, notwithstanding limited receiver coverage. This includes mapping movements in receiver gaps, which is an important development upon heuristic methods (15). By leveraging knowledge of the movement process plus the observations, we localised simulated individuals to within 0.4 % of the lake area. The potential for Bayesian techniques to further refine array design is evident from this analysis (45). We would expect gridded array designs, which minimise spaces where fish can remain without detection over prolonged periods, to refine our results. At regional scales, we estimated residency for simulated tracks with a median error and precision (standard error) below 1 %. This is a marked improvement upon heuristic analyses (28).

Our real-world analysis reveals patterns of movement and regional residency in nearly 100 lake trout. Building on previous work (28, 46), we provide robust confirmation of spawning site segregation, with north- and south-tagged fish returning to distinct spawning locations in fall but overlapping during other seasons. There is some evidence that north-tagged fish may exhibit elevated straying and lower spawning site fidelity, given their wider occurrence distribution during fall. However, broader distributions are also related to data quality, as uncertainty in individuals’ locations grows during detection gaps. This may explain higher residency estimates in regions such as Northeast Arm compared to previous analyses (28, 35, 46). Apparent hotspots in bays without receivers may also be linked to this effect, as they provide a refuge for particles during detection gaps. This highlights the importance of uncertainty quantification and domain expertise for interpretation and downstream ecological analyses (13, 16). While our maps correctly represent the model’s knowledge of an individual’s state, given available data and model assumptions, in situations where the data are not very informative the influence of those assumptions (especially the movement model) becomes dominating and it is up to practitioners to evaluate which of the possible movements are most plausible in reality (16).

Our outputs provide new opportunities to advance ecological understanding of the drivers of movement patterns. A key question for future work concerns the individual-level mechanisms that drive population-level differences in movement patterns. For example, in Lake Champlain, whether north-tagged fish individually exhibit wider movements (broader occurrence distributions) or occupy distinct, local areas (narrower, non-overlapping distributions), accounting for uncertainty (represented by the smoothing distributions). State-space modelling can help to move acoustic telemetry studies beyond pattern description to address ecological questions of this kind, which are relevant to many aquatic species (47), while contributing to broader scientific debates, such as the role of specialisation and generalisation in ecology (48).

The current work has broad management implications (49). In Lake Champlain, our results provide valuable baseline information on movement patterns for management, particularly in relation to future changes such as the planned cessation of stocking (50). During fall, the spatial segregation between north- and south-tagged fish provides further evidence for the co-existence of largely distinct stocks, which has implications for understanding the long-term prospects of a recovering population (28, 46). By advancing knowledge of movements, we can inform close-kin mark-recapture models of survivorship and other parameters crucial to management (50). Identification of unexpected, ecologically important movements is also valuable (51). For example, our results strengthen evidence of rare movements into the Northeast Arm during summer by north-tagged fish detected in this region. This is surprising considering expected environmental conditions (low dissolved oxygen levels) and advances our understanding of potential refuges during suboptimal conditions (52). Further investigation of these movements may also shed light on prey dynamics, which could supersede environmental conditions. By strengthening acoustic telemetry analyses, we hope to strengthen the links between acoustic telemetry and management in this way in Lake Champlain and beyond (49).

The particle algorithms developed here are one of several inference approaches (16). Their strengths include flexibility and scalability. The flexibility comes from the use of weighted samples (particles) to approximate probability distributions via forward simulation and likelihood evaluation plus resampling, without the requirement for gradients or closed-form analytical solutions (53). Consequently, particle algorithms can accommodate non-linear, non-Gaussian movement and observation models, including the land-truncated correlated random walk movement model we implemented and our bespoke, distance-decay detection probability function that requires a line of sight between transmitter and receiver. During inference, the recursive approximation of the marginal distribution *f*(***s***_*t*_ | ***y***_1:*T*_) is sufficient for mapping space use but simpler than sampling trajectories from the joint distribution *f*(***s***_1:*T*_ | ***y***_1:*T*_), as attempted by some alternative methods (25). This facilitates the analysis of big, high-dimensional datasets, which increasingly incorporate hundreds of fish of multiple species in systems around the world (27).

A limitation is particle degeneracy (53). This occurs when a few particles acquire most of the weight, which can lead to poor approximations and convergence failures. Degeneracy is akin to the evolutionary process of genetic drift (54) and mainly linked to sequential resampling of particles without conditioning on subsequent observations. This is exacerbated by limited prior knowledge for model specification, which makes inference harder. We attribute convergence failures in this study to this effect. For example, our movement model imperfectly represented switches between site-restricted behaviour and directional, longer distance movements. We observed specific issues in the south of the study area, where individuals sometimes passed through receiver barriers without detection. The bathymetry in this region is complex and we hypothesise areas with reduced detection probability that facilitate fish passage without detection. Targeted work to fill these data gaps identified by modelling should support refinement of our analyses in future.

Another limitation concerns the estimation of static parameters. For computational efficiency, we focused inference on location and defined parameters in the movement and observation models *a priori* using domain knowledge, complementary datasets and literature (16). In particle algorithms, parameter estimation is possible but computationally expensive (55). Our approach facilitates big data analysis but leads to an underestimation of uncertainty. Further research is needed to develop approaches, such as Hamiltonian Monte Carlo, for efficient joint estimation of latent states and parameters (56). The nature of acoustic telemetry systems makes this a challenging, but rewarding, problem for future research.

## 4. Materials and methods

### 4.1 Study system

Lake trout are large, freshwater, benthopelagic fish found across North America (20, 57). Growing up to 1.5 m in size over a 50-year lifespan, they principally inhabit deep, oligotrophic lakes, where they feed on plankton, invertebrates and fish and support recreational and commercial fisheries (20, 57).

In Lake Champlain (Fig. 1), a lake trout population has been restored after decades of management (58). The lake spans the borders of Vermont, New York and Quebec and is approximately 193 km long and 20 km wide. The depth averages 20 m (but reaches 122 m in its deepest point). For this study, we distinguish seven regions in the lake (Fig. 1).

Between 2013–17, an acoustic telemetry system was deployed (28, 46). From 2013–14, 93 fish were tagged in two sites (a northern and southern site) with 147 dB V13 acoustic transmitters, programmed with a nominal transmission interval of 120 s (Table S1). Concurrently, Vemco VR2W receivers were deployed at 31 stations. There were 153 receiver deployments during the study (Table S2). For this paper, data were sourced from refs. (28, 46). See Supporting Information §1 for additional considerations that informed analysis. For a summary of our workflow, see Supporting Information §2 or Fig. S1. For lake trout detections, see Fig. S2.

### 4.2. Model

#### Posterior

We formulated a discrete-time state-space model for lake trout movements (Fig. 2). Our task was to derive the joint probability distribution *f*(***s***_1:*T*_ | ***y***_1:*T*_) of each individual’s possible trajectories ***s***_1:*T*_ given the data ***y***_1:*T*_. By applying Bayes’ Theorem, we represent *f*(***s***_1:*T*_ | ***y***_1:*T*_) in terms of a movement process *f*(***s***_1:*T*_) and an observation process *f*(***y***_1:*T*_ | ***s***_1:*T*_)

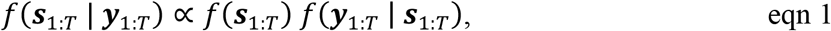

where the time steps *t* ∈ {1, 2, …, *T*} are spaced every two minutes (the nominal transmission delay).

We formulated models for the movement and observation processes, leveraging information in the literature, complementary datasets and domain expertise. All models were refined via model validation. For a summary of the datasets we analysed to inform model development, see Supporting Information §3 and Table S3. For details, see Supporting Information §4.

#### Prior

We modelled *f*(***s***_1:*T*_) as a discrete-time Markovian walk

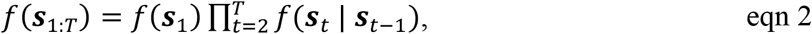

where *f*(***s***_1_) represents the probability distribution for the individual’s initial state and and *f*(***s***_*t*_ | ***s***_*t*−1_) represents the probability density of moving from ***s***_*t*−1_ → ***s***_*t*_. The state ***s***_1_ = (*s*_*x*,1_, *s*_*y*,1_, *ϕ*_1_) comprises the initial location (*s*_*x*,1_, *s*_*y*,1_) and the heading *ϕ*_1._

To model *f*(***s***_1_), we used a Uniform distribution for the starting location and initial heading:

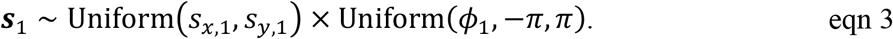

To model *f*(***s***_*t*_ | ***s***_*t*−1_), we used a two-dimensional correlated random walk. The location ***s***_*t*_ depends on the previous location ***s***_*t*−1_, a step length *d*_*t*_, the previous heading *ϕ*_*t*−1_ and a turning angle Δ*ϕ*_*t*_:

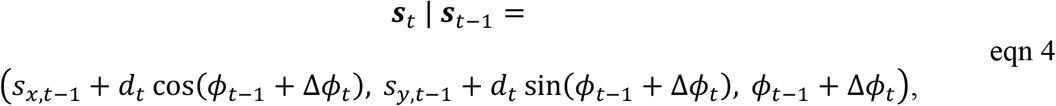

where *d* and Δ*ϕ* are independently distributed random variables, truncated by land and thermal habitat suitability in summer (bottom depth < 20 m).

We modelled step lengths *d*_*t*_ (metres per two minutes) with a truncated Gamma distribution

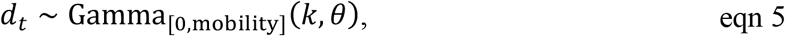

with location and scale parameters *k* = 3.25 and *θ* = 25, based on literature and our analysis of accelerometery measurements (20, 34) and fine-scale positioning data (21, 33). The distribution was truncated between zero and mobililty = 216 m, which defines the maximum moveable distance from *t* − 1 → t.

Turning angles Δ*ϕ* (radians) were modelled using a mixture distribution

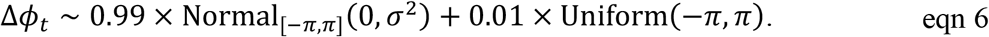

where *σ* = 0.4 radians. This model was informed by literature and our analyses of fine-scale positioning data (21, 33).

#### Likelihood

We modelled the observation process as a product over time steps

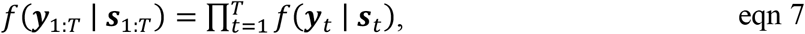

assuming independence. We denote the observation (non-detection, detection) at time *t* at receiver *k* as *y*_*t,k*_ ∈ {0, 1} and the receiver’s location by ***r***_*k*_ = (*x*_*k*_, *y*_*k*_). We modelled each observation *y*_*t,k*_ using the Bernoulli distribution

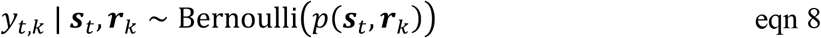

in which the probability of a detection is assumed to decline logistically with the Euclidean distance between the receiver and transmitter

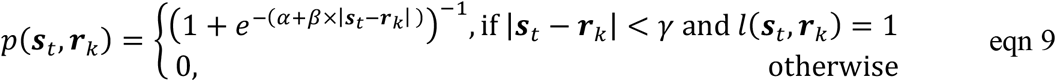

providing the distance is less than the maximum detection range and there is a line of sight between the location at ***s***_*t*_ and ***r***_*k*_. Line of sight *l*(***s***_*t*_, ***r***_*k*_) ∈ {0, 1} was approximated based on whether or not the midpoint between ***s***_*t*_ and ***r***_*k*_ was on land. Parameters were set to *α* ≈ 0.904, *β* ≈ −0.002 and *γ* = 7000 m. This model is based on literature and our analysis of in-situ (35, 46) and ex-situ (22) range-testing data.

For simulation sensitivity analyses, we also considered more restrictive and flexible parameterisations of the movement and acoustic observation models. For a summary of all parameterisations, see Table S4.

### 4.3. Inference

We performed inference using particle algorithms via the Patter.jl package, v.2.0.0 (43). This approach targets the marginal smoothing distribution *f*(***s***_*t*_ | ***y***_1:*T*_). The distribution is approximated with weighted particles derived via particle filtering and two-filter smoothing. Inference was performed for the latent states ***s***_*t*_, while leveraging prior information for static parameters. We analysed sensitivity to model parameterisation by repeating inference using restrictive and flexible model formulations. As diagnostics, we computed the area spanned by *f*(***s***_*t*_ | ***y***_1:*T*_) and effective sample sizes. Computation time was recorded on Debian machine with 2019 AMD EPYC 7742 server hardware. For details, see Supporting Information §5–7.

### 4.4. Simulation analysis

We first performed a simulation analysis to assess algorithm performance and sensitivity. We simulated 100 hypothetical trajectories and associated observations at each receiver station over a one-month period. For comparison to ref. (28), we retained the 96 trajectories associated with at least one detection and performed locational inference using our (a) data-generating, (b) restrictive and (c) flexible models. To assess performance and sensitivity, we visualised simulated trajectories versus occurrence distributions and computed mean absolute errors in residency for all individuals and lake regions. We also computed the Mean Weighted Occupancy Error (MOE), which summarises residency error over regions (28). For comparison, a heuristic (interpolation) method was included in this analysis which calculated residency from tracks interpolated between receivers (28). This analysis provided high-level validation of the routines plus quantitative assessments of method performance and sensitivity to parameter misspecification. For details, see Supporting Information §7.1.

### 4.5. Real-world analysis

Real-world data were processed following ref. (28). For analysis, we considered a timeline from 1^st^ December 2014 (after which time all individuals were tagged) until 31^st^ May 2017 (Fig. S2). For each individual, we performed locational inference every two minutes over the entire timeline (657,360 time steps). Inference was parallelised over batches of approximately monthly duration. For the two groups of north- and south-tagged lake trout, we collated seasonal occurrence distributions and residency by region, accounting for individual survivorship (50). For details, see Supporting Information §7.2. For the computational workflow, see Supporting Information §7.3.

## Author contributions

EL and MHF conceptualised the study. MHF provided the lake trout data. EL formulated the models, with substantial inputs from MHF, AS and CA. EL wrote the code, with additions from MHF, and analysed the data, with substantial inputs from MHF, AS and CA. EL wrote the first draft of the manuscript, with substantial contributions from MHF. CA, AS and SB advised on the mathematics. All authors (EL, MHF, AS, SB, JB, RAB, JT and CA) contributed to review and editing and approved publication.

## Acknowledgments

We thank Pinheiro et al. (46) and Futia et al. (28) for making the lake trout data available. Thanks to Paul Blanchfield and Tazi Rodrigues for collating and sharing their accelerometery data from lake trout (20), which informed our movement model, and comments on the manuscript. Connor Reeve shared model outputs for the conversion of accelerometer data into swim speeds (34). Fine-scale positioning data were sourced from Binder et al. (21) and Futia et al. (33). Natalie Klinard shared detection probability data (22), which informed our acoustic observation model. Additional detection probability data were provided by Futia and Marsden (35). Thanks to Helen Moor for additional support.

## Data availability

The Lake Champlain lake trout datasets (28) are archived on Zenodo (https://doi.org/10.5281/zenodo.17904694). Code is available on GitHub (https://github.com/edwardlavender/patter-champlain) and archived on Zenodo (https://doi.org/10.5281/zenodo.20498818).

