## Supporting Materials for "Big data analysis of animal movements in aquatic ecosystems with acoustic telemetry"


| **Contents** | **Pages** |
|---|---|
| **Supporting information text** | **2** |
| **1.** Study system | 2 |
| **2.** Workflow | 3 |
| **3.** Supporting datasets | 4 |
| **3.1.** Movement datasets | 4 |
| **3.2.** Observation datasets | 5 |
| **4.** State-space model | 5 |
| **4.1.** Movement model | 5 |
| **4.1.1.** Step length | 6 |
| **4.1.1.1.** Literature | 6 |
| **4.1.1.2.** Data analysis | 7 |
| **4.1.1.3.** Synthesis | 8 |
| **4.1.2.** Turning angle | 9 |
| **4.1.2.1.** Literature | 9 |
| **4.1.2.2.** Data analysis | 10 |
| **4.1.2.3.** Synthesis | 11 |
| **4.2.** Observation model | 11 |
| **4.2.1.** Literature | 11 |
| **4.2.2.** Data analysis | 12 |
| **4.2.3.** Synthesis | 13 |
| **5.** Inference | 14 |
| **5.1.** Locational inference | 15 |
| **5.2.** Estimation of occurrence and residency | 16 |
| **6.** Validation exercise | 16 |
| **7.** Analyses | 18 |
| **7.1.** Simulation analyses | 18 |
| **7.2.** Real-world analyses | 18 |
| **7.3.** Computational workflow | 20 |
| **Figures** | **22** |
| **Fig. S1**. A summary of the workflow. | 22 |
| **Fig. S2**. Detection time series. | 24 |
| **Fig. S3**. Simulation analysis: patterns of space use. | 25 |
| **Fig. S4**. Simulation analysis: residency patterns. | 26 |
| **Fig. S5**. Simulation analysis: residency patterns averaged over regions. | 27 |
| **Fig. S6**. Simulation analysis: diagnostics. | 28 |
| **Fig. S7**. Real-world analysis: lake trout residency in Lake Champlain. | 29 |
| **Fig. S8.** Real-world analysis: diagnostics, following Fig. S6. | 30 |
| **Tables** | **31** |
| **Table S1**. A summary of tagged fish. | 31 |
| **Table S2**. A summary of receiver deployments. | 34 |
| **Table S3**. A summary of ancillary datasets. | 40 |

| **Contents** | **Pages** |
|---|---|
| **Table S4**. A summary of parameter values. | 41 |
| **SI References** | **42** |

**Supporting information text**

### 1. Study system

**Geography.** Lake Champlain is one of the largest lakes in North America. There are five main basins (Missisquoi Bay, Northeast Arm, Malletts Bay, Main Lake and South Lake). The northern basins are separated by islands and causeways that form hydrological barriers (while permitting fish movement). For management purposes, the Main Lake is divided into three regions (North, Central and South). The lake flows north from the Champlain Canal in the south to the Richelieu River in the north, which leads to the St. Lawrence River. Locks and dams in the Champlain Canal and Richelieu River, respectively, heavily restrict fish movement (1). For the purpose of this study, we therefore treat Lake Champlain as a closed system. Using data from ref. (2), we represent the study system on a raster, with a grid resolution of 200 by 200 m (107,625 cells).

**Lake trout.** Lake trout (*Salvelinus namaycush*) are native to Lake Champlain but were extirpated by the 1900s (1). Initial reintroductions occurred in the 1950s and an annual lake trout stocking program began in the 1970s, re-establishing their population. Annual survival estimates have been variable over time and across life stages, but recent studies estimate adult annual survival probabilities (denoted $h$) for the stocked population of approximately 90 % (3, 4). We take the median value of $h = 0.93$ derived from close-kin mark recapture (4) as the best available estimate of annual adult survival probability for our analysis (see §7.2). Stocked lake trout were observed reproducing naturally at multiple nearshore sites throughout the Main Lake by the 2000s (5), but the population remained entirely supported by stocked fish until the 2010s when natural recruitment was first observed (6).

**Environmental conditions.** Within Lake Champlain, water temperatures encountered by lake trout vary from near zero to approximately 12 °C (7). Thermal stratification occurs in summer and, by August, the thermocline can reach depths of 20–30 m (8). During the stratified period, lake trout move into colder, deeper water and occupy the hypolimnion near the bottom of the lake (7, 9). In shallow areas above the thermocline, temperatures generally exceed lake trout temperature tolerances, which have an optimum temperature range of approximately 8–12 °C (9, 10). Lake trout can temporarily occupy habitat with temperatures exceeding 12 °C (10), but such excursions are exceptionally rare for lake trout studied in Lake Champlain (7, 9). During fall, the lake becomes isothermal and adult lake trout migrate to shallower habitat nearshore for spawning, and can remain there through spring (7, 9). Near-bottom current speeds are typically around 0.1–0.2 $ms^{-1}$, but can be up to 0.55 $ms^{-1}$ (11, 12). However, environmental conditions vary between basins.

**Data collection.** Data on the movements of lake trout in Lake Champlain were sourced from refs. (13, 14). Ninety-three fish (45 adult males and 48 adult females) were captured and tagged at two spawning locations in the study system during fall in 2013 and 2014. For details, see ref. (13), Fig. 1 and Table S1. Captured individuals ranged from 0.53–0.82 (median = 0.67) m in total length.

All individuals were tagged with Vemco (now Innovasea) V13 acoustic transmitters, operating at 69 kHz with a power output of 147 dB re 1 μPa at 1 m and a nominal delay of 120 s (Table S2). Transmitter weight (9.4 g) was less than 0.5 % the total body weight of individual lake trout (minimum weight: 2,160 g), limiting tag burden and the risk of tag expulsion or post-release mortality (15). Estimates for the latter are generally on the order of approximately 10

% (15–18). This shaped our data processing strategy (see below). Transmitter battery life was approximately four years (1583 days) and encompassed the full duration of our study. All individuals were detected. The raw dataset included 1,735,137 detections.

Receiver deployments (Vemco VR2Ws) began in fall 2013 (14). In summer 2014, additional receivers were deployed to expand receiver coverage. Thereafter, array design remained consistent through until spring 2017, after which time the array was rearranged to focus on a single location during summer 2017 (19). For the purpose of this study, we focus on the period from the start of winter 2014 to the end of spring 2017. All receivers were vertically suspended 2 m above bottom. Receiver depth ranged from 3.4–47.5 (median = 11.9) m.

**Data processing.** Data were processed following ref. (14). Key data processing steps included:

A. **Time correction.** Detection data were time-corrected for clock drift using VUE (20).
B. **False detection filter.** Detections were filtered to exclude potentially false detections flagged by the short-interval criterion (21).
C. **Mortality filters.** Fish presumed dead on the basis of regular detections at a single receiver until the end of the recorded time were excluded.

### 2. Workflow

To reconstruct lake trout movements in the Lake Champlain study system (§1), we followed a three-stage workflow (Fig. S1).

A. **Model development.** First, we undertook research to inform the development of a Bayesian state-space model for lake trout movements. The state-space model comprised a movement model, based on step lengths and turning angles, and an acoustic observation (detection probability) model. To inform model development, we sourced specific datasets (§3) with information on lake trout movements (§3.1) and detection probability (§3.2). We also reviewed key literature (§4.1.1.1, §4.1.2.1 and §4.2.1) and undertook data analyses (§4.1.1.2, §4.1.2.2 and §4.2.2).

B. **Model formulation.** Building on Stage A, we formulated a Bayesian state-space model (§4.1.1.3, §4.1.2.3 and §4.2.3). We formulated three versions of each (step length, turning angle and detection probability) sub-model, including a best, restrictive and flexible parameterisation (to analyse sensitivity). Model formulation was refined by a model validation exercise (§5–6). We took 100 real-world time series and performed inference. By analysing diagnostics, we identified specific issues in earlier model versions and iteratively refined the model.

C. **Inference.** We performed inference for the state-space model to estimate the locations of tagged individuals through time (§5). This was performed for two analyses (§7). First, we conducted a simulation-based analysis (§7.1). We simulated movement trajectories and acoustic observations for 100 hypothetical individuals and compared simulated patterns to those reconstructed from the observations. This analysis was used to examine the performance and sensitivity of the inference approach. Second, we conducted the real-world analysis in which we analysed real acoustic observations from lake trout (§7.2).

All data processing and analysis were implemented in `R`, version 4.5.x (22) and `Julia`, version 1.11.2 (23).

### 3. Supporting datasets

To inform our model, we sourced additional datasets with information on (a) lake trout movements and (b) detection probability in Lake Champlain and related systems. For a summary, see Table S3.

#### 3.1. Movement datasets

We sourced three datasets on lake trout movement properties (speeds, turning angles) for analysis.

**Alexie Accelerometry.** The Alexie Accelerometry dataset comprises 632,501 time-stamped accelerometer measurements from 14 lake trout. The data were collected by Blanchfield et al. (24) in a Vemco Positioning System (VPS) in Lake Alexie, northwestern Canada, from June 2013–14. Lake Alexie is a small (4 $km^2$) oligotrophic lake. The maximum depth is 32 m. Current speeds are presumed to be low. Of the lake trout systems studied by Blanchfield and colleagues, this is the most similar to Lake Champlain (Blanchfield, personal communication). The size (total length) range of tagged individuals was 0.52–0.61 (median = 0.58) m. Accelerometer tags were programmed with a 40 second sampling interval for which an average acceleration measurement was reported. We used these data, together with a swim speed calibration model (25), to learn about the distribution of swim speeds in lake trout (§4.1.1.2).

**Drummond Island VPS.** Lake trout positioning data were sourced from a VPS study at Drummond Island, Lake Huron, USA (26). The array comprised up to 140 VR2W acoustic receivers, covering 19–23 $km^2$ of the Drummond Island Refuge. The refuge includes multiple shoals and rocky reefs used by lake trout for spawning (27, 28). The acoustic tags were programmed to transmit signals at random intervals every 50–130 s. Data were collected in August–December in 2013 and over a similar period in 2014. Observations comprised timestamped individual position estimates with unitless horizontal position error (HPE) measurements. Over 75% of positions in 2013 and 2014 occurred during October and November (i.e., months associated with spawning behaviour). We estimated approximate horizontal position errors in metres for each fish position via calibration with sync tags of known position and position error. For our analysis of swim speeds (§4.1.1.2) and turning angles (§4.1.2.2), we focused on positions that had less than 20 m of estimated error based on HPE units, spaced less than 180 s apart (i.e., double the average transmission interval). These criteria retained 85 % of data while excluding time steps with longer durations during which time movements were uncertain. The processed dataset included 132 individuals and 583,939 positions. The total length range was 0.56–0.89 (median = 0.69) m.

**Thunder Bay VPS.** We also sourced lake trout positioning data from a VPS array deployed in 2021–2024 in Thunder Bay, Lake Huron (29). This system incorporated 40 acoustic receivers, including VR2Tx (2021–2023) and NexTrak R1 technology (2024). Receivers covered approximately 1.2 $km^2$ and were deployed around historic and recently constructed reefs during fall of each year. The study system is shallow (< 10 m) and was designed to target spawning movements. Acoustic tags were programmed to transmit signals at random intervals every 240–360 s. Data comprised position and error estimates, as in Drummond Island, but with better positioning accuracy. Therefore, we focused on positions with errors below 5 m and less than 300 s apart (the nominal transmission interval). These criteria retained 80 % of positions. The processed dataset included 66 individuals and 412,883 positions. The total length range was 0.54–1.15 (median = 0.70) m.

#### 3.2. Observation datasets

Range testing data for passive acoustic telemetry were obtained from two studies (7, 30).

**Champlain Range Test.** Range testing data were sourced from one study in Lake Champlain (7). This study conducted 172 range tests using V9 (146 and 151 dB) and V13 (152 dB) tags from 2021–2. For each test, a high-frequency (7 or 15 s) transmitter was suspended by a surface buoy and positioned approximately 3–5 m above lake bottom at a set distance from a receiver for approximately 30 minutes. We call each deployment an 'event'. Tests were conducted in nine locations broadly covering a similar area to receivers in the present study. Two of the nine locations aligned with receivers in the current study. The raw data comprised 10,652 timestamped detections of transmitters (in known locations) at receivers (in known locations). We translated this information into a data table of the observed versus expected number of detections at each receiver per event, which we used to model detection probability (§4.2.2).

**Ontario Range Test.** We also sourced a more extensive range testing dataset from eastern Lake Ontario, Canada (30). Range tests were performed in a deep underwater valley in the lake and included eight range-testing tags, programmed with a nominal delay of 1800 s, and 90 acoustic receivers deployed below the thermocline. We focused on data from three V9 (145 dB) tags (deployed at 11–50 m depth) and one V13 (153 dB) tag (deployed at 50 m). The power output range of these tags envelopes technology used in our Lake Champlain system (§1). The raw data comprised 293,786 detections at 67 receivers, collected from October 2015–May 2016. Following ref. (30), we assembled a data table of the observed/expected number of detections per day for analysis. Following the terminology above, we consider each day an 'event'.

### 4. State-space model

We formulated a Bayesian state-space model for lake trout movements in the Lake Champlain passive acoustic telemetry system (see Main Text Methods and Fig. 2). Given the sparsity of the receiver array (relative to the lake size), we based our model parameterisation on available datasets, domain expertise and literature and focused model inference on the latent locations. This section explains how we derived the movement and observation components of the state-space model.

#### 4.1. Movement model

In our Bayesian state-space model, the movement model is a prior. We formulated the prior as a correlated random walk with a time-invariant distribution of step lengths and turning angles, truncated by land and thermal habitat suitability in summer (see below). This was designed to be (a) simple, (b) generalisable and (c) sufficiently flexible to capture the distribution of possible movements while (d) preventing impossible movements.

In all analyses, the prior was truncated by land using our raster of the study area (see §1 and §5 for implementation details). In summer, we additionally truncated the prior by thermal habitat suitability, since seasonal changes in lake temperature are a primary factor driving lake trout habitat use (31). We assumed movements into areas less than 20 m depth were impossible. This is the shallow bound for the thermocline depth (§1). To implement this threshold, we sourced a bathymetry point dataset from ref. (32). We rasterized the dataset onto our study area raster by triangulated interpolation, using the `interpNear` function in the `terra` package, version 1.8-86 (33). Movements into cells above the 20 m threshold were assigned zero probability. Including this additional information in the model improved inference in summer when detections were sparse (§6). This filter retained some habitats in the Northeast Arm and Malletts Bay of suitable depth (temperature) but likely insufficient dissolved oxygen (34). However, as dissolved oxygen measurements are only available at specific point locations, we did not include oxygen suitability in our model.

We recognise that in real-world ecosystems, individual movements vary with the time of day, food web structure, seasonality and other variables (35). However, these effects vary among systems and are not fully understood, so we did not attempt to include them in our prior. To parameterise the prior, we examined literature studies of lake trout and other fish species, analysed supporting datasets and leveraged domain expertise.

##### 4.1.1. Step length

We modelled the distribution of step lengths using a truncated Gamma distribution informed by literature (§4.1.1.1) and analyses of accelerometry measurements, fine-scale positioning and acoustic detections (§4.1.1.2).

###### 4.1.1.1. Literature

There is a wide literature on teleost locomotion (36–38). Here, we summarise results from specific studies we leveraged to inform a model of step lengths (movement speeds) in lake trout.

**Cruz-Font et al. (39)** studied lake trout swim speeds from performance in swim tunnels and field observations from three boreal lakes in the Experimental Lakes Area in northwest Ontario. In swim-tunnel experiments, they increased swim speed by 0.035 $ms^{-1}$ for ~5 min until fish reached fatigue. Slow speeds were defined as <0.3 $m^{-1}$ or <0.5 fork lengths per second ($FLs^{-1}$). Fast speeds were defined as >0.57 $ms^{-1}$ (>1 $FLs^{-1}$). At these speeds, tag drag frequently exceeded swimming capacity and fish exhibited erratic bursting movements. Individuals hit fatigue at speeds of 0.8–0.95 $ms^{-1}$ (>1.5 $FLs^{-1}$). The authors also estimated swimming speeds in wild fish, accounting for movement in three dimensions, using fine-scale radio-acoustic (VRAP) positional telemetry. Swimming speeds in the field ranged from approximately 0 to > 0.3 $ms^{-1}$. By comparing results from short- (~25 s) and long-delay (4 min) transmitters, they showed that linear movements (especially slower ones) generally underestimated distance travelled by two times (suggesting potential speeds up to 0.9 $ms^{-1}$ in the field, in line with laboratory results).

**Blanchfield et al. (24)** studied trout speed using fine-scale positional telemetry (VPS) and accelerometer tags in Lake Alexie (§3.1). Average daily array speeds (derived from positional telemetry) varied from ~0.05–0.2 $ms^{-1}$. Average daily accelerometer speeds were higher, varying from approximately 0.19–0.44 $ms^{-1}$. However, these summary statistics underestimate the true variability in swim speeds (see §4.1.1.2).

**Reeve et al. (25)** expanded previous work using swim tunnel respirometry (39) to evaluate swimming performance and associated metabolic costs in lake trout (§3.1). Most swim speeds ranged between 1.0 and 1.5 total body lengths per second ($BLs^{-1}$). Speeds increased with fish size and ambient temperature. Maximum observed swim speeds approached 2.5 $BLs^{-1}$ for the highest temperature treatment (15 °C).

**Blanchfield et al. (40)** evaluated lake trout swim speed during spawning and non-spawning periods in response to boat presence. Average swim speeds during 1-hour tracking periods were during non-spawning periods were approximately double calculated speeds observed during spawning periods. While this comparison was not the focus of the authors, it demonstrates that assessments of lake trout movement patterns on spawning grounds during spawning periods may be biased towards slower movements compared to movements during other seasons, which is relevant for our consideration of the Drummond Island and Thunder Bay VPS datasets (§3.1).

**Pike and Burman (41)** determined fine-scale, three-dimensional fish movements in a controlled environment and used those data to simulate biologically-plausible movement patterns using three-spined stickleback (*Gasterosteus aculeatus*) as a model organism. They observed times with limited movement as well as intermittent burst movements during 60-minute observation periods. While the biology of three-spined stickleback differs from lake trout, the authors found that a gamma distribution was a suitable model for variable swim speeds and we take this as a starting point for our model of step lengths in lake trout.

##### 4.1.1.2. Data analysis

For more detailed information on lake trout swim speeds, we analysed our detection dataset (§1) plus the Alexie Accelerometry, Drummond Island VPS and Thunder Bay VPS datasets (§3.1).

**Detection analysis.** For the Lake Champlain system, we computed the speeds required to produce the observed pattern of sequential detections at receivers, following refs. (42, 43). We found that movement speeds up to 0.91 $ms^{-1}$ are required to produce the observed pattern of detections, if detections were recorded when individuals were close to receivers (the most likely scenario). However, for movements between the outer edges of the detection range (§4.2), speeds up to 1.53 $ms^{-1}$ are required to reproduce detections. These results are broadly consistent for different (random) transmission intervals and detection ranges (§4.2).

**Alexie Accelerometry analysis.** We reconstructed an 'observed' distribution of swimming speeds (step lengths), using the raw acceleration measurements provided by Blanchfield et al. (24). To convert acceleration into swim speed, we used Reeve et al.'s best speed ~ acceleration model (25). This model relates speed ($S$) in body lengths per second ($BLs^{-1}$) to an acceleration $A$ measurement $i$ from an individual $j$ by the equation:

$$\log_{10}(S_{ij}) \sim \beta_0 + \beta_1 \log_{10}(A_{ij}) + u_j + \varepsilon_i \qquad \text{eqn S1}$$

where $\beta_0$ and $\beta_1$ are fixed effects, $u_j \sim N(0, \sigma^2_{\text{id}})$ is an individual-specific random effect and $\varepsilon_i \sim N(0, \sigma^2_{\text{resid}})$ is the residual error.

The parameter estimates for this model derived by Reeve et al. (25) were:

- Fixed effect maximum likelihood estimates: $\hat{\beta}_0 = 0.15353, \hat{\beta}_1 = 0.36788$
- Fixed effect standard errors: $\tilde{\beta}_0 = 0.01529, \tilde{\beta}_1 = 0.02338$
- Correlation between fixed effects: $\rho = 0.224$
- Standard deviation among individuals: $\sigma^2_{\text{id}} = 0.05566$
- Residual standard error: $\sigma^2_{\text{resid}} = 0.04809^2$

For each measurement, we simulated 250 swimming speeds (indexed by $k$) from equation S1:

$$\log_{10}\left(S_{ij}^{(k)}\right) \sim \beta_0^{(k)} + \beta_1 \log_{10}\left(A_{ij}\right) + u_j^{(k)} + \varepsilon_i^{(k)}, \qquad \text{eqn S2}$$

where each pair of fixed effects was sampled from the bivariate normal distribution

$$\begin{bmatrix} \beta_0^{(k)} \\ \beta_1^{(k)} \end{bmatrix} \sim N\left(\begin{bmatrix} \hat{\beta}_0 \\ \hat{\beta}_1 \end{bmatrix}, \begin{bmatrix} \tilde{\beta}_0^2 & \rho\tilde{\beta}_0\tilde{\beta}_1 \\ \rho\tilde{\beta}_0\tilde{\beta}_1 & \tilde{\beta}_1^2 \end{bmatrix}\right), \qquad \text{eqn S3}$$

each random effect $u_j^{(k)}$ was sampled from a normal distribution

$$u_j^{(k)} \sim N(0, 0.05566^2), \qquad \text{eqn S4}$$

and residual error $\varepsilon_i^{(k)}$ terms were sampled similarly:

$$\varepsilon_i^{(k)} \sim N(0, 0.04809^2). \qquad \text{eqn S5}$$

Aggregated across all measurements, this produced a distribution of swimming speeds, accounting for all modelled sources of uncertainty (Fig. 2). The mean speed was 0.85 $BLs^{-1}$; the standard deviation was 0.32 $BLs^{-1}$ and the 99$^{th}$ percentile (an indicator of the maximum movement speed, excluding outliers) was 1.98 $BLs^{-1}$. We confirmed the distribution was robust to (a) autocorrelation in the dataset by repeating the analysis with thinned datasets in which with only one measurement retained for each individual per (i) hour, (ii) 2 hours and (iii) 12 hours and (b) data structure, by repeating the analysis with random subsamples of the data derived without replacement containing (i) 25 %, (ii) 50 % and (iii) 75 % of observations.

For the purpose of the present study, we translated the aggregated distribution of swimming speeds ($BLs^{-1}$) to a distribution of step lengths ($ms^{-1}$ or m per two minutes) for the (i) smallest and (ii) largest individuals in our study (Fig. 2). For the smallest fish, the summary statistics of this analysis are 0.47 (mean), 0.17 (standard deviation) and 1.09 (99$^{th}$ percentile) $ms^{-1}$. For the largest fish, the same statistics are 0.70, 0.26 and 1.62 $ms^{-1}$ (or 84, 31 and 195 m per 2-min) respectively. Our model for the distribution of step lengths was derived to envelop these features exhibited by the observations.

**Drummond Island VPS analysis.** We calculated swim speeds for the VPS datasets from estimated positions (Fig. 2). We excluded the fastest 5% of movements for each fish to remove erroneous estimates associated with poor positioning (up to approximately 20 $ms^{-1}$). The dataset included 205,923 estimated speeds. Across all individuals, the median swim speed was 0.06 $ms^{-1}$ (median absolute deviation: 0.07 $ms^{-1}$). Individual medians ranged from 0.03–0.71 $ms^{-1}$. Maximum swim speeds ranged from 0.18–0.88 $ms^{-1}$ (median = 0.5 $ms^{-1}$).

**Thunder Bay VPS analysis.** In the Thunder Bay VPS dataset, we estimated 217,871 speeds (Fig. 2). The median swim speed was also 0.06 $ms^{-1}$ (median absolute deviation: 0.06 $ms^{-1}$). Individual medians ranged from 0.02–0.28 $ms^{-1}$. Maximum swim speeds ranged from 0.15–0.58 $ms^{-1}$ (median = 0.36 $ms^{-1}$).

##### 4.1.1.3. Synthesis

Building on the work above, we derived three models for the distribution of step lengths (Fig. 2):

**A. Best model**

$$d_t \sim \text{Gamma}_{[0,216]}(3.25, 25.00) \qquad \text{eqn S6}$$

**B. Restrictive model**

$$d_t \sim \text{Gamma}_{[0,162]}(4.33, 16.88) \quad \text{eqn S7}$$

**C. Flexible model**

$$d_t \sim \text{Gamma}_{[0,270]}(2.60, 35.17) \quad \text{eqn S8}$$

All models were subject to boundary conditions; that is, land and thermal habitat suitability in summer (see §4.1).

The best model exhibits the following characteristics:

- **A. Lower bound.** We assume lake trout are constantly moving, with movements faster than 0.05 m/s increasing in probability.
- **B. Central region.** We assume a broad peak of moderate probability movements between 0.1–1.2 $ms^{-1}$. This encompasses the lower speeds estimated from the two VPS datasets and aligns with the Alexie Accelerometry analysis.
- **C. Upper bound.** The upper bound was chosen to envelop the speeds we observed from positional telemetry (which likely underestimate swimming speeds) and the results of the Alexie Accelerometry analysis. For largest fish (0.82 m length), a maximum swimming speed of 1.5 $BLs^{-1}$ or 1.25 $ms^{-1}$ seems plausible (§4.1.1.1). In areas of high near-bottom current flow, potential movement speeds may be up to 0.55 $ms^{-1}$ higher (§1). Thus, we consider a maximum possible movement speed for the trout tagged in this study of 1.8 $ms^{-1}$, which approximately aligns with the swimming speeds implied by our Alexie Accelerometry analyses. In the model, such speeds are possible, but unlikely. The exact threshold is not critical, providing it is large enough (44). Movement speeds beyond 1.8 $ms^{-1}$ are assumed impossible.

The restrictive and flexible models represent a 25 % reduction/expansion in flexibility. These models represent suitable checkpoints for the sensitivity analyses, as we were able to adjust the turning angle (§4.1.2.3) and detection probability (§4.2.3) models by the same amount while keeping behaviour within sensible bounds. However, they do not cover the full range of uncertainty.

#### 4.1.2. Turning angle

As for step lengths, we derived a model for the distribution of turning angles based on literature (§4.1.2.1) and data analyses (§4.1.2.2).

##### 4.1.2.1. Literature

As a first step to inform the turning angle model, we reviewed information in the literature on movement tortuosity in lake trout and related species.

**Hawkins et al. (45)** reviewed biomechanics associated with fish swimming and provided a general assessment of turning behaviour. Turning is an important component of fish locomotion. Importantly, trout and other species that use their bodies to turn typically make wider turns (i.e., greater turning radii) than species that primarily use their pectoral fins for turning. However, many factors can influence turn angle. In particular, periods of faster movement are often more directed, as fish transit from one location to another. This review suggests that capturing the capacity for both turning and directed movement is important for modelling fish behaviour (see §4.1.2.3).

**Blanchfield et al. (40)** evaluated lake trout behaviour and movement patterns in response to manual tracking to determine if fish react and alter their behaviour in response to boat activity. The researchers evaluated fish depth, swim speed and path predictability with and without boat traffic. Under both conditions, lake trout had similar turning behaviour. However, substantial differences in turn angle were observed between spawning and non-spawning periods. When occupying spawning habitat (i.e., shallow reefs), lake trout tend to have a much higher average turn angle per hour (approximately 1.9 radians) compared to non-spawning periods (approximately 1.3 radians). This indicates more direct movements associated with dispersal during non-spawning periods compared to more localised movements (e.g., circling a spawning reef) during the spawning period. This is an important consideration the inclusion of the Drummond Island and Thunder Bay VPS datasets in our analysis (see §4.1.2.3).

**Christiansen et al. (46)** studied the three-dimensional movement patterns of the marine, mesopelagic fish *Maurolicus muelleri* using a surface-facing, moored echosounder. Turning behaviours were assessed based on average tortuosity calculated across the duration of individual fish tracks (18 seconds to 12 minutes). Tortuosity followed a right-skewed distribution with most tracks having relatively straight lines regardless of time of day. Smaller turning angles were observed during migratory periods and were associated with directed movement. While this species is very different from lake trout, this analysis adds to the evidence that fish can exhibit periods of directed movements, which we need to consider in our model.

**Pike & Burman (41)** evaluated fish turn angles using observations and simulations for three-spined stickleback (§4.1.1.1). Most fish typically maintained forward motion with minimal change between steps (i.e., straight movement) and occasional turns at large angles. As noted above, smaller turns were also linked to more directed movements in this species. However, the frequency of turns and range of turn intensity was variable among individuals. These results suggest that some flexibility in models of turn angles is required.

##### 4.1.2.2. Data analysis

To inform our turning angle model, we analysed positions from the Drummond Island and Thunder Bay VPS datasets (Fig. 2). We recognise that these datasets are limited by the site-restricted nature of the VPS arrays, which were designed to track fish within specific areas of interest (i.e., reefs) and timeframes. We therefore expected the data to capture turns more frequently than typical, as over broader spatiotemporal scales larger movements between disparate habitats may occur (14). However, they provide a quantitative baseline for model development.

**Drummond Island VPS.** We obtained 244,797 turn angles from the Drummond Island VPS dataset (Fig. 2). Angles (radians) were estimated as the angle generated among three consecutive positions (i.e., start, middle and end position) by individual fish. We focused on estimates for which the time between the two steps used to calculate the turn angle (start position to middle position, middle position to end position) was less than six minutes. Over all individuals, the distribution of turning angles exhibited a main peak at 0 radians (directional movement) with lower peaks at $\pm \pi$ radians (180° turns) and a broad spread in-between, in line with our expectations. The standard deviation in turning angles was 1.7 radians. Within the domain of the VPS array, these results suggest generally correlated movements but with periods of randomness and changes in direction.

**Thunder Bay VPS.** An additional 19,182 angles were calculated from the Thunder Bay VPS dataset (Fig. 2). Due to the longer tag delay for this second dataset (§3.1), we focused on angles computed over a time interval below 9 min (the smallest cutoff that still provided data). The results were similar to the Drummond Island VPS dataset, with a median turning angle of 0.1 radians and a standard deviation of 1.8 radians.

##### 4.1.2.3. Synthesis

Building on the work above, we initially formulated a relatively flexible model for turning angles:

$$\Delta\phi_t \sim \text{Normal}(0, 1.8^2). \quad \text{eqn S9}$$

However, during model validation, we observed that this model was unable to generate longer, more directed movements between disparate receivers (§6). After further evaluation, we therefore derived the following three models (Fig. 2):

A. **Best model**

$$\Delta\phi_t \sim 0.99 \times \text{Normal}_{[-\pi,\pi]}(0, 0.4^2) + 0.01 \times \text{Uniform}(-\pi, \pi) \quad \text{eqn S10}$$

B. **Restrictive model**

$$\Delta\phi_t \sim 0.99 \times \text{Normal}_{[-\pi,\pi]}(0, 0.3^2) + 0.01 \times \text{Uniform}(-\pi, \pi) \quad \text{eqn S11}$$

C. **Flexible model**

$$\Delta\phi_t \sim 0.99 \times \text{Normal}_{[-\pi,\pi]}(0, 0.5^2) + 0.01 \times \text{Uniform}(-\pi, \pi) \quad \text{eqn S12}$$

As for the step length model (§4.1.1.3), all models were subject to boundary conditions.

The models were designed to capture directed movement (with a normal distribution) and uncorrelated changes in direction (with a uniform distribution). The parameterisation of the best model was chosen to capture both site-restricted and large-scale movements in the validation analysis (§6). The restrictive and flexible models represent a 25 % increase/decrease in flexibility and define pragmatic checkpoints at which to examine sensitivity in simulations. We recognise that these models do not capture the full range of uncertainty but permit both correlated and uncorrelated movements to varying degrees. Together with the step-length models, they form our priors. These were updated by the likelihood to infer individuals' locations (§4.2 and §5).

#### 4.2. Observation model

The observation model provides the link between the unknown locations of a tagged animal and the observations, which comprised detections and non-detections at receivers (47). Typically, we model detections with a Bernoulli ('coin toss') distribution, where the probability of a detection declines with distance from a receiver, though many variables can affect detection probability (48). To inform our model, we reviewed literature (§4.2.1) and analysed selected datasets (§4.2.2).

##### 4.2.1. Literature

Many studies have analysed detection probability in passive acoustic telemetry systems (48). We reviewed selected examples from Lake Champlain and related study systems.

**Pinheiro et al. (13)** conducted a 48 hour range test of one receiver with 147 dB V13 transmitters in August 2014. The receiver was located in the northern section of the Main Lake basin, Lake Champlain. Individual transmitters were deployed near the bottom at four distances (50, 100, 250, and 500 m) from the receiver for the 48-hour period. At distances of 50 and 100 m from the receiver, approximately 75 % of transmissions were detected. At 250 m, 70 % of transmissions were detected. By 500 m from the receiver, one third of transmissions were detected.

**Futia & Marsden (7)** evaluated spatial and temporal variability in detection range for three of the primary basins in Lake Champlain: Main Lake, Northeast Arm and Malletts Bay (§3.2). Multiple range tests were conducted during spring, summer and fall. Each test used a V9 (146 dB or 151 dB) and/or V13 (152 dB) transmitter deployed in fixed locations for approximately 30 minutes. Nine receivers were used for the tests (VR2W and VR2Tx technology operating at 69 kHz). Detection distances varied by transmitter. Higher power transmitters were detected at greater distances. Detection distances also varied by location and season, but no consistent patterns were identified. The largest distance with 50 % detection efficiency was approximately 1,450 m and observed in the Main Lake during fall.

**Klinard et al. (30)** studied detection efficiency in Lake Ontario (§3.2). They deployed an array of 90 VR2W 69 kHz receivers in deep water (below the thermocline during stratification). Range tests were performed using multiple transmitters (V9, 145 dB; V13, 153 dB; V16, 158 dB) at fixed locations and depths. Transmitters with higher power and positioned deeper tended to have greater detection efficiency than weaker and shallow transmitters. However, shallow transmitters had the greatest maximum detection ranges (6.4 km for the V9 transmitter and 9.3 km for the V16 transmitter).

**Brownscombe et al. (49)** quantified variation in detection ranges in an array of 60–89 69 kHz VR2W and VR2Tx receivers deployed in the Florida Keys. Short-term (2 min) range tests were conducted at a range of distances from each sentinel receiver using a V13 transmitter placed 1 m below the surface. Distances of 77–364 m marked the 50 % detection efficiency threshold. By distances of 95–451 m, detection efficiency was reduced to 5 %. These results add to the evidence base that detection probability is strongly distance dependent across a wide range of systems and environmental conditions. Alongside distance, they found that detection ranges were generally higher for receivers with deeper water and less complex environments (low rugosity), which may be important in Lake Champlain (see Main Text Discussion).

**Long et al. (50)** studied detection efficiency in an array of 20 VR2W 69 kHz receivers deployed in a shallow-water (< 10 m) marine environment: Wellfleet Harbor, Massachusetts, USA. Receivers were deployed throughout the harbour. A preliminary mobile range test using a towed 152 dB V13 transmitter was conducted as well as three separate stationary range tests, lasting one to seven days, with transmitters positioned at multiple fixed locations near receivers. Detection efficiency declined with distance from receivers but, even close to receivers, detection probability was imperfect. The maximum detection range was 1,628 m. Alongside distance, the importance of other variables such as line of sight between receivers and transmitters was recognised, which is a relevant consideration for Lake Champlain (see §4.2.3).

##### 4.2.2. Data analysis

**Data preparation.** Following the literature review, we sourced the Champlain and Ontario Range Test datasets, from Futia et al. (14) and Klinard et al. (30), for analysis (§3.2). We considered each study (F, K) and tag power (dB) combination as a separate dataset. This produced five datasets for analysis, labelled 'F146', 'F151', 'F152', 'K145' and 'K153'.

**Modelling**. For each dataset, we analysed detection efficiency by comparing the total of detections per event ($i$) of each transmitter ($j$) at each receiver ($k$), in relation to the expected number of detections, as a function of distance between the transmitter location at $\boldsymbol{s}_j$ and receiver location $\boldsymbol{r}_k$. We opted for a Binomial Generalised Linear Model (GLM) of the form

$$y_{ijk} \mid \boldsymbol{s}_j, \boldsymbol{r}_k \sim \text{Binomial}\left(n_{ijk}, p\left(\boldsymbol{s}_{ij}, \boldsymbol{r}_k\right)\right) \quad \text{eqn S13}$$

where $n_{ijk}$ is the expected number of detections and detection probability $p\left(\boldsymbol{s}_{ij}, \boldsymbol{r}_k\right)$ declines as a logistic function of the distance between the transmitter and receiver:

$$p\left(\boldsymbol{s}_{ij}, \boldsymbol{r}_k\right) = \left(1 + e^{-(\alpha + \beta \times |\boldsymbol{s}_{ij} - \boldsymbol{r}_k|)}\right)^{-1}. \quad \text{eqn S14}$$

We opted for the binomial GLM because it is a simple model that is broadly generalisable across systems (48). We recognise other variables affect detection probability (§4.2.1), but their effects cannot be incorporated in models in the absence of sufficient data. However, we visually assessed the suitability of the GLMs by visual comparison with GAMs in which detection probability was modelled as a smooth function $g$ of distance, i.e.,

$$p\left(\boldsymbol{s}_{ij}, \boldsymbol{r}_k\right) = g\left(\left|\boldsymbol{s}_{ij} - \boldsymbol{r}_k\right|\right). \quad \text{eqn S15}$$

We used a thin-plate regression spline for the smoothing basis, with a basis dimension of five. Models were fitted using the `stats` (22) and `mgcv` (51) packages.

**Results.** In all analyses, there was a clear, negative relationship between detection probability and distance (Fig. 2). Detection probability declined more steeply in the Champlain Range Tests than the Ontario Range Tests. Within each system, detection probability functions were steepest for the lowest powered tags. Estimated 50 % detection ranges ranged from 233 m (F146) to 1484 (K153). Both GLM and GAM models produced similarly shaped functions. There was no evidence for non-linearity in the detection probability function that was not captured by the GLMs. For the Champlain Range Test dataset, observed maximum detection ranges were 1064 m (146 dB), 2306 m (151 dB) and 2320 m (152 dB). For the Ontario Range Test, maximum detection ranges were 6364 m (145 dB) and 8176 m (153 dB).

##### 4.2.3. Synthesis

We formulated a logistic, distance-decaying detection probability model. Initially, we derived a model based principally on the Ontario Range Test dataset. The model took the form

$$p(\boldsymbol{s}_t, \boldsymbol{r}_k) = \begin{cases} \left(1 + e^{-(1.886 - 0.002\,|\boldsymbol{s}_t - \boldsymbol{r}_k|)}\right)^{-1}, \text{if } |\boldsymbol{s}_t - \boldsymbol{r}_k| < 8000 \\ 0, \qquad \text{otherwise} \end{cases} \quad \text{eqn S16}$$

where $\boldsymbol{s}_t$ includes the transmitter location at time $t$, $\boldsymbol{r}_k$ is receiver $k$'s location and 8000 m is a truncation parameter that defines a maximum detection range. This defines the threshold beyond which detection probability changes from negligible (1.6 x $10^{-5}$ at 8000 m) to impossible (0 at 8001 m). Setting a threshold can improve computational efficiency during inference with particle algorithms, though the precise value has a limited bearing on locational inference providing it is large enough (§5). The term $p(\boldsymbol{s}_t, \boldsymbol{r}_k)$ is the probability for a detection at $\boldsymbol{r}_k$ given the transmission from the location in $\boldsymbol{s}_t$. (For inference, note that the term $p(\boldsymbol{s}_t, \boldsymbol{r}_k)$ is used as a weight, which is later normalised.)

However, the model validation phase showed this model was overly flexible (§6). That is, the model effectively blocked certain movements through receiver 'gates' which occurred without detection in the south of study area, where the geometry of the lake is highly constrained. We therefore revised the workflow, through examination of the more limited Champlain Range Test dataset and revision of our assumptions.

This led to the following models (Fig. 2):

**A. Best model**

$$p(\boldsymbol{s}_t, \boldsymbol{r}_k) = \begin{cases} \left(1 + e^{-(0.9039 - 0.0021\,|\boldsymbol{s}_t - \boldsymbol{r}_k|)}\right)^{-1}, \text{if } |\boldsymbol{s}_t - \boldsymbol{r}_k| < 7000 \text{ and } l(\boldsymbol{s}_t, \boldsymbol{r}_k) = 1 \\ 0, \quad \text{otherwise} \end{cases} \qquad \text{eqn S17}$$

**B. Restrictive model**

$$p(\boldsymbol{s}_t, \boldsymbol{r}_k) = \begin{cases} \left(1 + e^{-(0.6779 - 0.0026\,|\boldsymbol{s}_t - \boldsymbol{r}_k|)}\right)^{-1}, \text{if } |\boldsymbol{s}_t - \boldsymbol{r}_k| < 5250 \text{ and } l(\boldsymbol{s}_t, \boldsymbol{r}_k) = 1 \\ 0, \quad \text{otherwise} \end{cases} \qquad \text{eqn S18}$$

**C. Flexible model**

$$p(\boldsymbol{s}_t, \boldsymbol{r}_k) = \begin{cases} \left(1 + e^{-(1.1299 - 0.0016\,|\boldsymbol{s}_t - \boldsymbol{r}_k|)}\right)^{-1}, \text{if } |\boldsymbol{s}_t - \boldsymbol{r}_k| < 8750 \text{ and } l(\boldsymbol{s}_t, \boldsymbol{r}_k) = 1 \\ 0, \quad \text{otherwise} \end{cases} \qquad \text{eqn S19}$$

The best model lies between the detection probability functions derived for the F146 and F151 datasets. This model is more flexible than expected for a 147 dB tag from the Champlain Range Test dataset alone but accounts for results from the larger Ontario Range Test dataset while remaining less flexible than our initial model. We set the maximum detection range to 7000 m. This lies within the region defined by the K145 (6364 m) and K153 (8176 m) datasets and aligns with our reading of the literature (§4.2.1).

We accounted for line-of-sight $l(\boldsymbol{s}_t, \boldsymbol{r}_k) \in \{0, 1\}$ following review of the behaviour of the model around Grand Isle in the northern part of the study system (§6). For computational efficiency, $l(\boldsymbol{s}_t, \boldsymbol{r}_k)$ was computed approximately by checking whether the mid-point between the location at $\boldsymbol{s}_t$ and $\boldsymbol{r}_k$ was in water (1) or on land (0) according to our raster of the study area. The criterion $l(\boldsymbol{s}_t, \boldsymbol{r}_k) = 0$ assigns a detection probability of zero to transmissions blocked by land. However, this does not account for causeways which were not resolved on our map, where we expect some signal obstruction. There were seven receiver stations that could have been influenced by this, but six of those receivers were in basins where lake trout positions were less common (i.e., shallow water) so we do not judge this to be a serious limitation.

The restricted and flexible models represent and 25 % steepening/shallowing of the detection probability function respectively, in line with adjustments to the movement model (§4.1.1.3 and §4.1.2.3).

### 5. Inference

### 5.1. Locational inference

To estimate individual locations through time, we used the particle filtering–smoothing algorithm implemented by the `Patter.jl` package (52). This procedure targets the marginal distribution $f(\boldsymbol{s}_t \mid \boldsymbol{y}_{1:T})$ of the individual's location $\boldsymbol{s}_t$ at each time $t$, accounting for all observations $\boldsymbol{y}_{1:T}$. This target is sufficient for mapping space use and estimating residency, and cheaper to approximate than the joint distribution $f(\boldsymbol{s}_{1:T} \mid \boldsymbol{y}_{1:T})$ of the individual's trajectories $\boldsymbol{s}_{1:T}$ given $\boldsymbol{y}_{1:T}$ (47). This implementation of this algorithm was a two-step process.

**Particle filter.** The particle filter approximates the 'partial' marginal distribution $f(\boldsymbol{s}_t \mid \boldsymbol{y}_{1:t})$ of the individual's states $\boldsymbol{s}_t$, given all preceding data $\boldsymbol{y}_{1:t}$ (53). The key idea is to represent the unknown locations of the individual probabilistically, with a 'cloud' of particles. The main steps are as follows.

- **Initialisation.** We initialised the algorithm by sampling particles uniformly within the region of the study area compatible with the initial observations. (This is equivalent to uniform sampling over the whole study area, once the likelihood with respect to the first time point is applied, but more efficient and the default behaviour in `Patter.jl`.) A recursive simulation is then run in which particles are sampled from a movement model (the prediction) and weighted by the data (the update).
- **Movement.** During the recursion, we sampled particles from our correlated random walk model (equations S6–8 and S10–12). To implement the truncation by land and summertime thermal habitat suitability, we repeatedly sampled movements for each particle until a valid movement was identified. Particles for which a valid movement was not sampled after 1000 attempts were killed.
- **Acoustic observations.** Particles were weighted in line with their compatibility with the acoustic observations (detections, non-detections at each operational receiver) using the acoustic observation model (equations S17–19).
- **Acoustic containers.** To improve algorithm convergence, we included an additional observation structure termed an 'acoustic container' (54). This defines the region within which an individual must be located at any time $t$ according to the receiver(s) at which it was next detected. We used this structure to kill particles which were incompatible with the location of future detection(s). For computational efficiency, we only implemented containers when they shrank to less than half the size of the study area (i.e., 44159.98 m); that is, if the time until the next detection was less than approximately 3.5 hours, according to our best mobility estimate (§4.1.1.3). This process can improve convergence in particle algorithms, which are otherwise blind to the future. (In the absence of acoustic containers, in a sparse receiver array, many particles do not reach the location of the next detection and are killed, which degrades the quality of the approximation.)
- **Resampling.** We used adaptive low variance resampling to resample particles when the Effective Sample Size ESS $\leq$ 1000 particles. To promote convergence, we also forced resampling at moments in time when the acoustic containers came into operation. The resampling process duplicates particles with large weights and eliminates particles with zero weight. The collection of surviving particles at each time step forms a heatmap of the animal's possible locations, approximating $f(\boldsymbol{s}_t \mid \boldsymbol{y}_{1:t})$.

The computational cost of this procedure grows linearly with the number of particles $N$ and time steps $T$; that is, $O(NT)$. Many particles can be required to compensate for particle degeneracy/death and to achieve adequate Effective Sample Sizes (ESS $\geq$ 500 is desirable to approximate a two-dimensional distribution). In simulations, we used 10,000 particles (§7.1). We revised this upwards to 50,000–100,000 particles for the real-world analysis (§7.2).

**Particle smoother.** To approximate $f(\boldsymbol{s}_t \mid \boldsymbol{y}_{1:T})$, the two-filter smoother was used (55). This entails running the particle filter forwards and backwards in time and then a smoothing operation that reweights particles in line with the probability density of moves between pairs of particles on the two filters. The computational cost of this operation increases with the square of the number of particles; that is $O(N^2T)$. Therefore, the number of particles that can be used for smoothing is more restricted than for filtering. This can problematic if there are relatively few valid routes between particle pairs the two filters. If there are no valid routes between the subset of retained particles, `Patter.jl` returns 50 % of particles from the forward filter and 50 % of particles from the backward filter, as proper smoothing is not possible (43). We used 1,500 particles for simulation analyses (§7.1) and 2,500 particles for real-world analyses (§7.2). We only analysed outputs for which proper smoothing was possible on more than 75 % of time steps (our convergence threshold).

Algorithms were implemented in single-threaded mode on a Debian server with 2019 AMD EPYC 7742 server hardware. Algorithm runs were parallelised over time series.

#### 5.2. Estimation of occurrence and residency

**Occurrence.** Using smoothed particles, for each time series we reconstructed the occurrence distribution and estimated residency in the seven regions of Lake Champlain (47). The occurrence distribution is a heatmap that defines the probability of observing the individual in a given location. Mathematically, the occurrence probability $P$ in each grid $A$ is defined as the time-averaged probability mass in that cell

$$P(A) = \frac{1}{T}\sum_{t=1}^{T}\int_{A\times S^1} f(\boldsymbol{s}_t \mid \boldsymbol{y}_{1:T})\, d\boldsymbol{s}_t, \qquad \text{eqn S20}$$

where $A$ denotes the area of interest and $S^1$ denotes the unit circle (i.e., we integrate out the headings).

Since we approximated $f(\boldsymbol{s}_t \mid \boldsymbol{y}_{1:T})$ with $N$ weighted particles, we computed the occurrence probability from the number of particles in each grid cell

$$P(A) = \frac{1}{T}\sum_{t=1}^{T}\sum_{i=1}^{N} I_A(\boldsymbol{s}_{i,t})\, w_{i,t}, \qquad \text{eqn S21}$$

where $\boldsymbol{s}_{i,t}$ includes the location of the $i^{\text{th}}$ particle at time $t$; $I_A$ is an indicator function that returns 1 if the location defined by $\boldsymbol{s}_{i,t}$ is in $A$ or 0 otherwise; and $w_{i,t}$ denotes the normalised weight.

**Residency.** Using the occurrence distribution, we estimated residency in each region of the lake. Residency is defined as the percentage Perc of time spent in each region $R$ and was computed as the sum of all occurrence probabilities in that region

$$\text{Perc}(R) = \sum_{A\in R} P(A) \times 100. \qquad \text{eqn S22}$$

### 6. Validation exercise

We fine-tuned the state-space model formulation and the inference settings with a validation analysis. In this analysis, we implemented the particle filter for 100 randomly selected real-world individual/month time series. We investigated the causes of convergence failures, which occur when none of the simulated particles are compatible with the observations, by visualising animated maps of particle movements (that is, $f(\boldsymbol{s}_t \mid \boldsymbol{y}_{1:t})$ snapshots) through time in relation to the observations. As this process was instructive, we describe some insights here for the benefit of readers developing particle algorithms in other study systems.

**Step lengths.** In an earlier version of this manuscript, we formulated the step length model based on literature, especially ref. (24), and the Drummond Island and Thunder Bay VPS datasets (§3.1). We found this model was too restrictive. We sought additional quantitative data on swim speeds in lake trout and obtained the Alexie Accelerometry and swim-tunnel calibration datasets for analysis (24, 25). This work suggested a broader range of swimming speeds and a lead to a refinement of the step length model.

**Turning angles.** We similarly found that an initial, weakly correlated turning angle model (equation S9) was limiting. This model was unable to capture movement between more distant receivers. (Animations showed particles spreading out too slowing to reach necessary receivers.) This led to a more correlated model, which was able to capture both site-restricted behaviour and more extensive movements between disparate sites. Within Lake Champlain, the more correlated model was initially computationally expensive, as by default `Patter.jl` truncates the movement model and each particle move that arrives on land is re-simulated 100,000 times until a valid move is reached (the `n_move` setting). We improved performance by incorporating a uniform distribution in the model for turning angles and adjusting the `n_move` setting to 1,000 times.

**Movement truncation.** Our revised movement model appeared to work reasonably well in fall, winter and spring when relatively regular detections help to 'correct' the prior. However, model behaviour was poorer in summer when detections were extremely sparse (Fig. S2). During this time, we observed particles moving into thermally inhospitable nearshore habitats without receivers. To mitigate this issue, we refined the model with knowledge of summertime thermal habitat suitability.

**Detection probability.** Our initial detection probability model (equation S16) was based principally on the extensive Ontario Range Test dataset, informed by literature (§3.2 and §4.2.1). While this model was generally compatible with the observations, it was overly flexible. In particular, in the south of the study area, we observed that the movement of particles between disparate receivers was effectively blocked by 'gates' of intermediate receivers which individuals sometimes passed through without detection (under the flexible model, passing through without detection was highly unlikely). We therefore incorporated the Champlain Range Test dataset and refined the detection probability model accordingly. After visualising particle animations, it also became clear that it was necessary to account for line of sight. This ensured sensible behaviour of the model around Grand Isle in the north of Lake Champlain, which in places is narrower than the maximum detection range.

**Geometry.** Even with a refined state-space model, we observed challenges in certain parts of the study area. For example, in the south of the study area we found that small particle populations (partially isolated by receiver gates) can be vulnerable to collapse: in narrow channels, it is hard to avoid movements onto land, which induce particle death. Mitigating strategies included boosting the number of particles and the number of attempts per particle used to generate a valid move, and reducing resampling. However, these strategies tended to facilitate inference for some time series but not others. Increased computational effort was also expensive.

On the basis of the above validation analysis, we settled on the aforementioned formulation of the state-space model and the inference settings. These settings achieved a reasonable balance between success and computational effort.

### 7. Analyses

#### 7.1. Simulation analyses

We conducted a simulation analysis to quantify the performance and sensitivity of the inference approach within the study area, following ref. (43).

**Simulation.** We simulated 100 trajectories using best movement model (equations S6 and S10). Each simulation was randomly initialised on our study area raster and continued at a time resolution of two minutes for one month. At each time step, we simulated acoustic observations at each of the 31 receiver stations using best observation model (equation S17). For comparison to ref. (14), we focused subsequent analysis on the 96 % of trajectories which were associated with at least one detection. We visualised each trajectory and computed residency in each of the seven regions of the lake.

**Inference.** Using the simulated observations, we reconstructed individual movements via particle filtering and smoothing (§5.1). To align with the real-world analysis, we did not include 'tagging' locations in this process, though this would refine inferences. As diagnostics, we examined convergence rates and ESS. Occurrence distributions were estimated on our study area raster (§5.2). Residency was estimated from the percentage of smoothed, resampled particles in each region. For comparison to ref. (14), we compared these residency estimates to those derived from the best-performing heuristic (interpolation) method identified by ref. (14), termed *Int*. This method calculates residency using tracks interpolated between receivers by linear and non-linear interpolation. Following ref. (14), we applied this approach using `interpolate_path` function in the `glatos` package (56).

**Analysis.** To analyse algorithm performance, we visually assessed the correspondence between simulated trajectories and estimated occurrence distributions. We computed the median area spanned by 95 % of the probability mass as a metric of spatial uncertainty. For residency, we computed the mean absolute error (ME) in simulated versus estimated residency in each region. We took the median absolute deviation in ME across trajectories as a measure of the precision with which we can expect to estimate residency for any one individual, given the properties of the study system and individual variation in movement patterns. Following ref. (14), we also computed Mean Weighted Occupancy Error (MOE) values as an average of the residency error over all regions. Repeating this analysis using mis-specified (restrictive, flexible) state-space model parameterisations provided an assessment of sensitivity under controlled conditions.

#### 7.2. Real-world analyses

**Data processing.** In the real-world analysis, we analysed movement patterns using the real passive acoustic telemetry dataset (§1). In total, 69 individuals were included in the analysis. Data were processed for analysis as described in §1. To align detections with the time resolution of inference procedure, we rounded detections to the nearest two-minutes. Duplicate detections (at the same receiver in the same two-minute period) were dropped.

**Inference.** We performed locational inference for each individual every two minutes for 657,360 time steps from the start to the end of the timeline (2014-12-01 00:00:00 until 2017-05-31 23:58:00). That is, we continued the inference process beyond the last detection for each individual, accounting for survival probabilities after this time during the collation of the results (see below). This approach avoids inappropriately biasing the results towards receiver locations (which would happen if we only modelled movements during the detection period for each individual). The inference process was handled in blocks (§7.3). For the particle filter, we used 50,000 particles. Filter runs that failed to converge were repeated with 100,000 particles. For the particle smoother, we used 2,500 particles. In line with the results of the simulation analysis and to minimise the number of computations, we only performed inference using our best model formulation for the real-world time series. We did not include the heuristic comparison (§7.1).

**Analysis.** We computed occurrence distributions and residency by region, following the simulation analysis, for each individual and season/year. Seasons were defined following ref. (14).

Using the above occurrence distributions, we computed the overall distribution for each season from the (normalised) sum of all modelled occurrence distributions

$$P(A) \propto \sum_{j=1}^{J} P_j(A)\, S_j, \qquad \text{eqn S23}$$

where $j$ indexes individual/year combinations and $P_j(A)$ is the probability that a given individual in a given season/year block was in grid cell $A$. This treats the maps from each individual as independent across years. The term $S_j$ denotes the probability that the individual was alive in that season/year block (see below).

Similarly, we computed the average residency in each region $R$ per season, accounting for survival probability $S_j$, as

$$\mathrm{Perc}(R) = \frac{\sum_{j=1}^{J} \mathrm{Perc}_j(R)\, S_j}{\sum_{j=1}^{J} S_j}, \qquad \text{eqn S24}$$

where $j$ indexes individual/year combinations and $\mathrm{Perc}_j(R)$ is estimated percentage of time that a given individual in a given season/year block spent in region $R$. Note that $\mathrm{Perc}(R)$ sums to 100 % over all $R$.

**Survivorship weights.** For both the occurrence and residency analyses, survival weights were defined as follows. For each individual, for all season/year blocks up to and including the last detection for that individual, $S_j = 1$. For individuals that were detected by later array deployments after the end date of this study (57), $S_j = 1$ for all subsequent blocks. For the remaining individuals, for blocks after their last detection we computed $S_j$ from a constant hazard model

$$S_j = h^{\tau_j}, \qquad \text{eqn S25}$$

where $S_j$ is the probability of the individual surviving $\tau_j$ years after its last detection and $0 < h < 1$ is the annual survival probability. Using the best available estimate of annual survival probability for lake trout in Lake Champlain (§1)

$$h = 0.93, \qquad \text{eqn S26}$$

we computed the survival probability of each individual for each block after the last detection as

$$S_j = 0.93^{\tau_j}. \qquad \text{eqn S27}$$

In practice, we computed $\tau_j$ as the (fractional) number of years from the date of the last detection to the midpoint of that season/year block.

#### 7.3. Computational workflow

To manage the computations for the real-world analysis, we considered three strategies. For the benefit of readers developing analyses in other systems, in this section we evaluate these options and summarise the workflow we adopted.

**Contiguous strategy.** An obvious strategy is to run, for each individual, the inference procedure from the start to the end of the timeline. That is, we run the filter forwards, backwards and then we implement smoothing. There are three limitations with this approach.

1. **Degeneracy.** Long time series are more susceptible to particle degeneracy and convergence failures.
2. **Storage.** Long time series can have significant memory or disk-space requirements. `Patter.jl` can handle memory requirements for long time series by periodically writing particles to disk. For our analysis, we have 657,360 time steps and 2,500 particles per time step, each of which contains three double-precision floating-point numbers: the (x, y) coordinates and the previous heading. Each double occupies 8 bytes. Since this information needs to be stored for both forward and backward runs and then the smoother, in our case this approach would have required up to 657,360 × 2500 × 3 × 3 × 8 / $1\times10^{-9}$ = 118 GB per time series (or 8,164 GB if we handled all 69 individuals simultaneously).
3. **Scalability.** Long time series runs limit the efficiency with which we can exploit High-Performance Computing (HPC) infrastructure for parallelisation. Long jobs are also penalised by HPC schedulers.

**Regular blocking.** A second option is to split each time series into regular blocks, such as seasons. For each block, the algorithms are run. After smoothing, outputs are summarised as needed and then particle files can be deleted. This approach mitigates issues regarding particle degeneracy, storage and scalability. However, the treatment of blocks as independent time series results in a loss of information: information outside the block of interest, which could help initialise the forward or backward filter, is not considered.

**Data-driven blocking.** We used a third option. We defined smaller blocks of monthly duration. We expanded each one-month block forwards and backwards in time by up to one month to ensure, where possible, that each block started and ended with a detection. This produced a set of 2070 blocks, up to approximately three months in duration. We treated all blocks as effectively independent: at the moment of a detection, particles cluster around the receiver(s) that recorded the detection, so little information is added by considering previous or later time steps. To reduce computational burden, we only implemented the algorithms for 1427 blocks with at least one detection in the surrounding blocks(s). These blocks were handled in parallel. For each block, we ran the filters and smoother. After smoothing, we removed the particle files for the filters, summarised smoothed particles and then removed the particle files for the smoother. Smoothed particles were summarised by counting the number of particles in each grid cell per time step. This reduced disk space requirements because the number of grid cells was generally much less than (and always less than or equal to) the number of particles. For the remaining blocks, which included all blocks more than one month after the block containing the last detection for each individual, we assumed a uniform distribution over study area (accounting for habitat suitability in summer). This assumption aligns with the long-run behaviour of the movement model. After the inference process, we collated the outputs for groups of blocks, which we term ‘chains’, into occurrence distributions and residency estimates (§7.2). For our analysis, each chain represents an individual/season/year grouping. Outputs from multiple chains were then collated to investigate overall patterns of space use and residency by season and tagging location.

### Figures

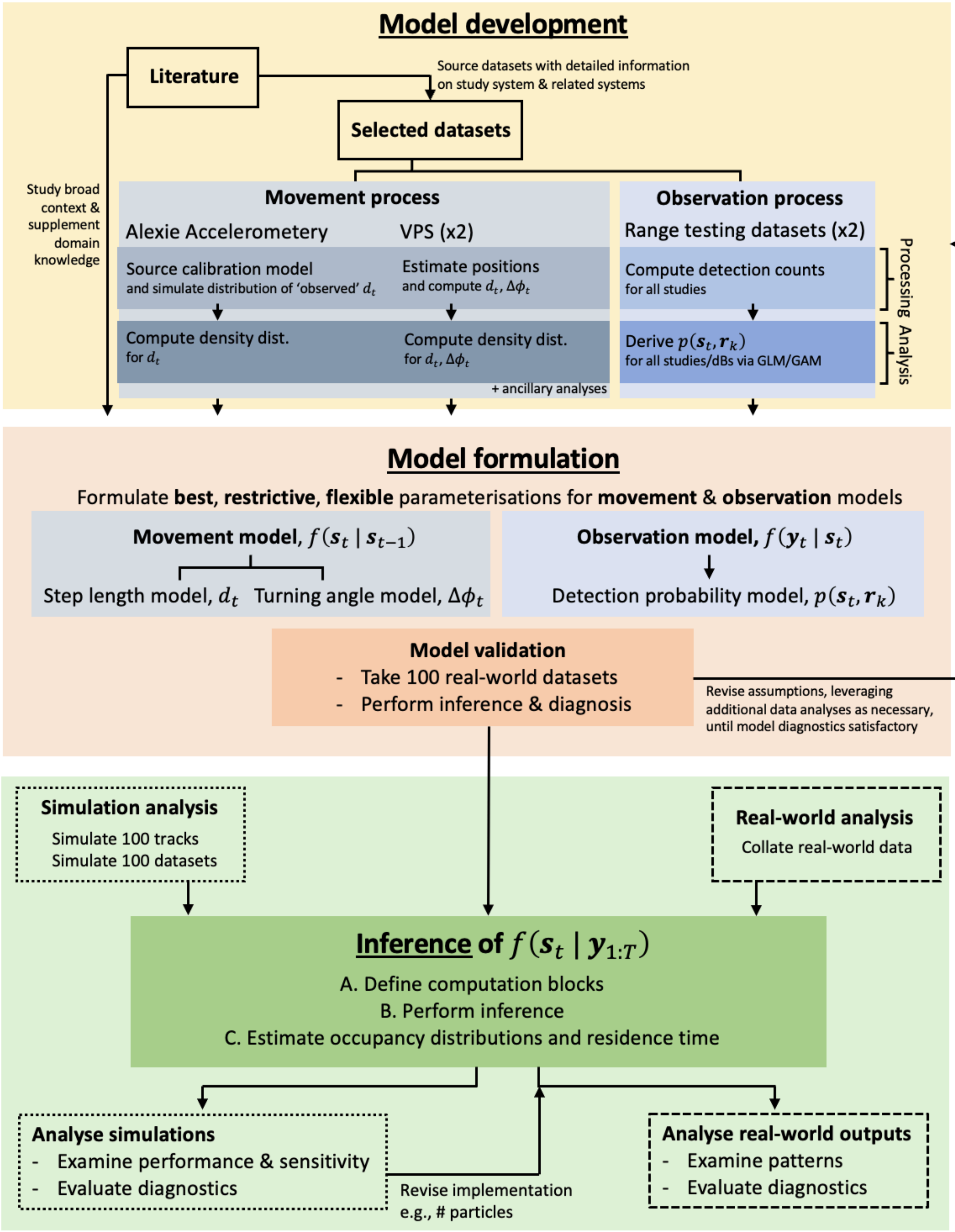
Model development
Literature
Source datasets with detailed information on study system & related systems
Selected datasets
Study broad context & supplement domain knowledge
Movement process
Alexie Accelerometery
VPS (x2)
Source calibration model and simulate distribution of 'observed' $d_t$
Estimate positions and compute $d_t$, $\Delta\phi_t$
Compute density dist. for $d_t$
Compute density dist. for $d_t$, $\Delta\phi_t$
+ ancillary analyses
Observation process
Range testing datasets (x2)
Compute detection counts for all studies
Derive $p(s_t, r_k)$ for all studies/dBs via GLM/GAM
Processing
Analysis
Model formulation
Formulate best, restrictive, flexible parameterisations for movement & observation models
Movement model, $f(s_t \mid s_{t-1})$
Step length model, $d_t$
Turning angle model, $\Delta\phi_t$
Observation model, $f(y_t \mid s_t)$
Detection probability model, $p(s_t, r_k)$
Model validation
- Take 100 real-world datasets
- Perform inference & diagnosis
Revise assumptions, leveraging additional data analyses as necessary, until model diagnostics satisfactory
Simulation analysis
Simulate 100 tracks
Simulate 100 datasets
Real-world analysis
Collate real-world data
Inference of $f(s_t \mid y_{1:T})$
A. Define computation blocks
B. Perform inference
C. Estimate occupancy distributions and residence time
Analyse simulations
- Examine performance & sensitivity
- Evaluate diagnostics
Revise implementation e.g., # particles
Analyse real-world outputs
- Examine patterns
- Evaluate diagnostics

**Fig. S1. A summary of the workflow.** To develop the state-space model for lake trout, we reviewed key literature and sourced specific datasets containing detailed information on the processes of interest. Leveraging literature, data analyses and domain knowledge, we formulated a Bayesian state-space model. We derived best, restrictive and flexible parameterisations (to analyse sensitivity). We iteratively refined these choices via a model validation step for a sample of 100 real-world datasets (individual/month time series). By studying the behaviour of the model for these time series, we were able to diagnose and resolve issues in the initial model formulation. For example, we found: (a) an initial movement model, based principally on ref. (24), was overly restrictive; (b) an initial turning angle distribution, informed by site-restricted VPS analyses (26), was incompatible with directed movements between disparate receivers; and (c) an initial detection probability model, based largely on ref. (30), in tandem with lake geometry, blocked certain observed movements through receiver 'gates' which occurred without detection. These observations led to refinement of the model, for example via the addition of the Alexie Accelerometry analysis (24, 25), until model diagnostics were satisfactory. Following initial model validation, we performed inference. We focused inference on the latent states, given best, restrictive or flexible settings for static parameters. We conducted simulation and real-world analyses. The simulation analysis was used to examine algorithm performance and sensitivity in a controlled setting and to inform the real-world analysis. For the real-world analysis, we examined movement patterns and diagnostics.

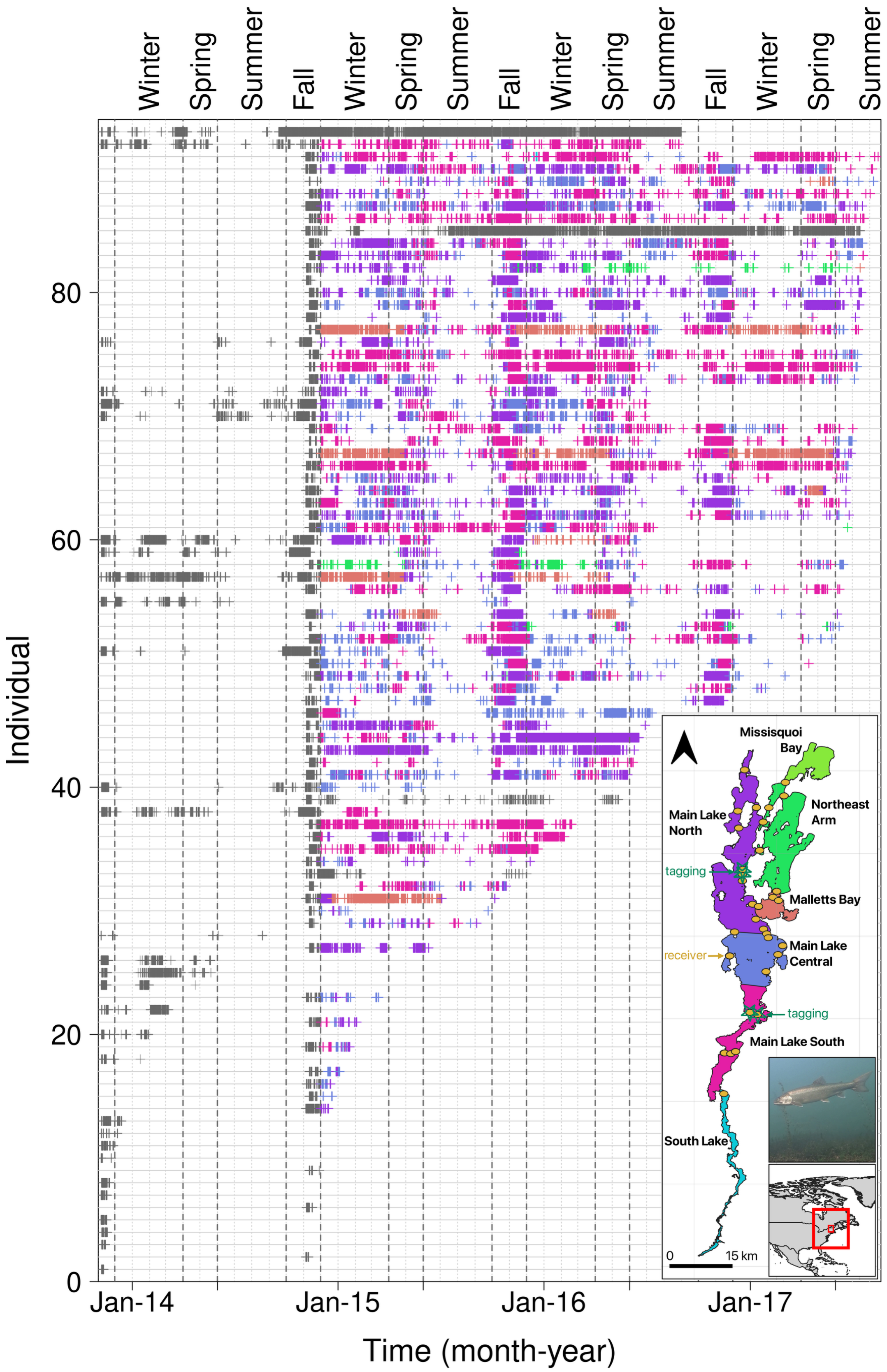


**Fig. S2. Detection time series.** Points mark detections of individuals, through time. Individuals are labelled following Table S1. Gridlines mark individuals/months. Points are coloured by receiver location (see inset). Time series that were excluded from the analysis are shown in grey.

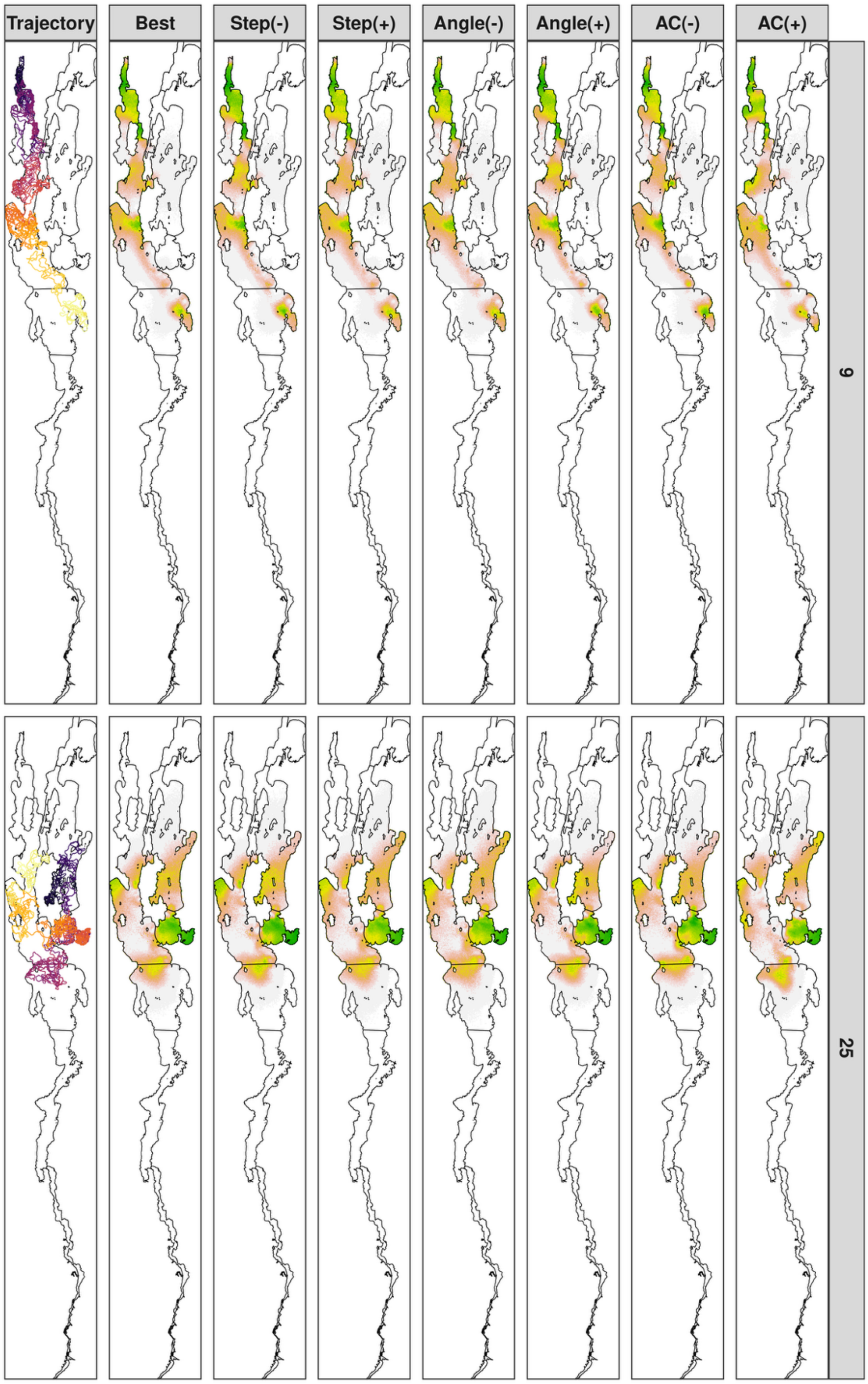


**Fig. S3. Simulation analysis: patterns of space use.** Each panel shows the simulated trajectory (first column) and the reconstructed occurrence distributions (columns two to eight) for a selected individual (row). In the main analysis, we reconstructed occurrence distributions using data-generating parameters (Best); in the sensitivity analysis, we reconstructed distributions using restrictive (-) and flexible (+) parameterisations of the step length (Step), turning angle (Angle) and acoustic observation (AC) models. Trajectories are coloured by time (black → yellow). On the occurrence distributions, colours follow Fig. 3 and show probability quantiles.

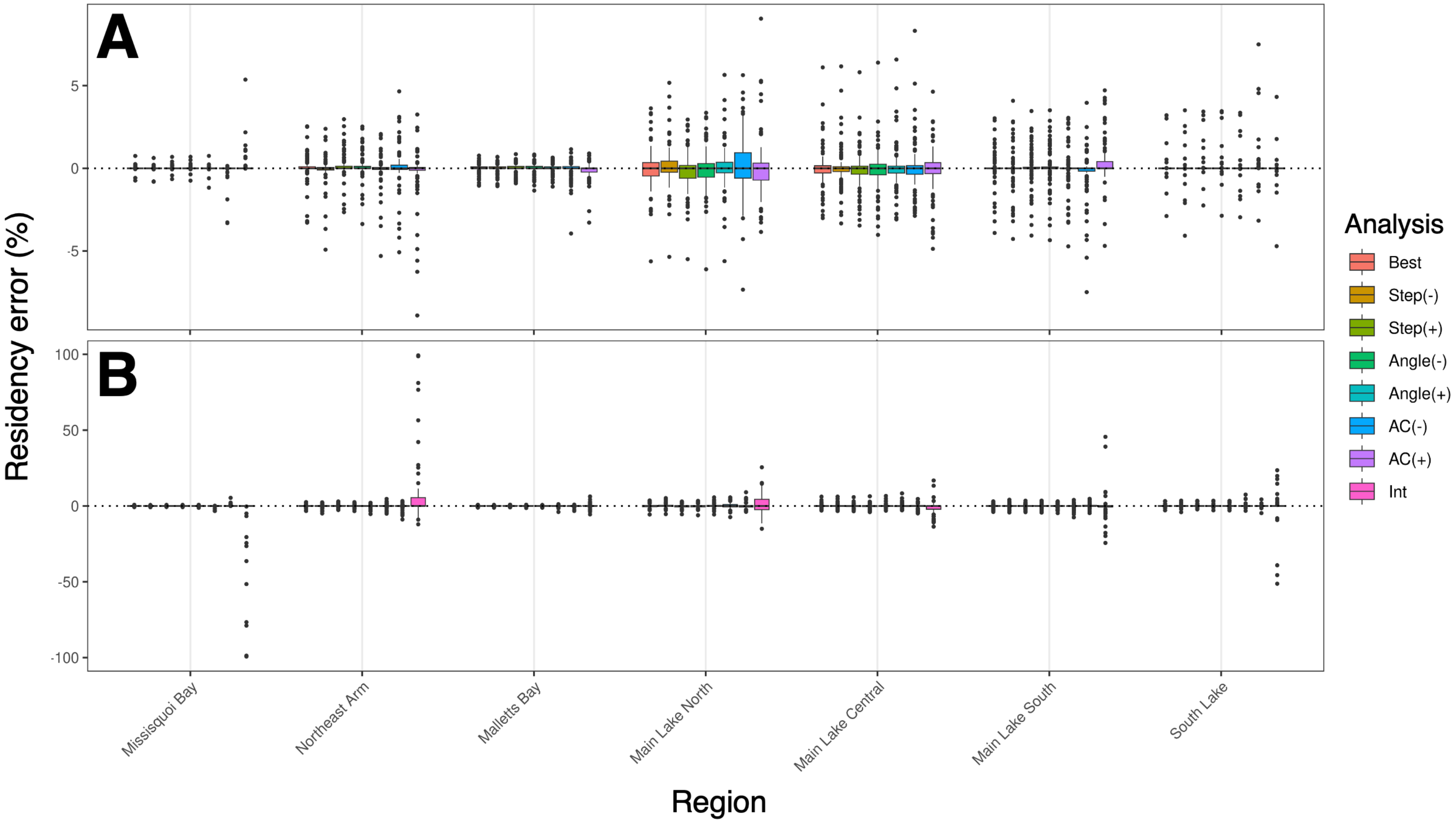


**Fig. S4. Simulation analysis: residency patterns. A** shows results from state-space modelling, following Fig. S3. **B** includes results from a heuristic (interpolation) method *Int*, which span a wider y-axis range and are included for comparison. Boxplots show the difference between the simulated and estimated percentage of time steps spent in a given region according to a selected analysis for 100 simulated individuals. The thick black line marks the median, the box edges mark the first ($Q_1$) and third ($Q_3$) quartiles and bar ends mark the range (excluding statistical outliers). Points mark statistical outliers (values $< Q_1 - 1.5 \times \text{IQR}$ or $> Q_3 + 1.5 \times \text{IQR}$, where IQR is the interquartile range). Box width is proportional to the number of algorithm runs that converged.

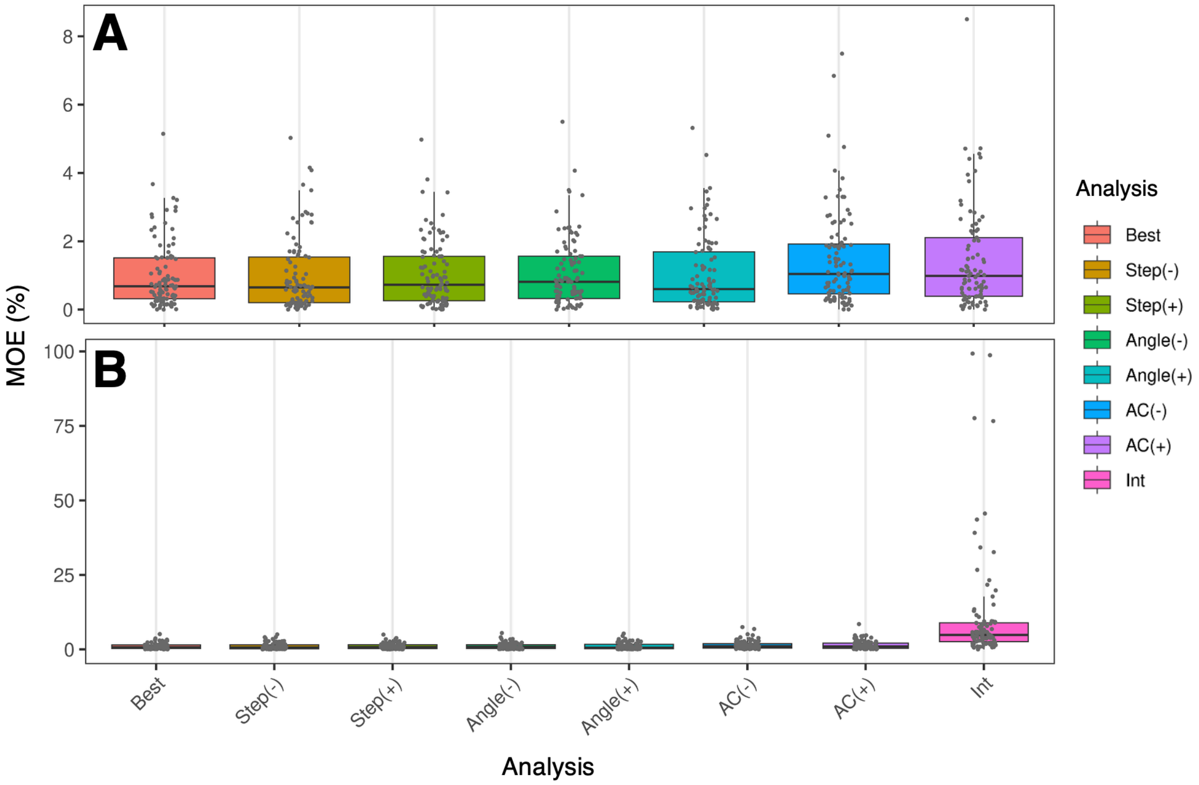


**Fig. S5. Simulation analysis: residency patterns averaged over regions.** Following Fig. S4, **A** shows results from state-space modelling and **B** includes results from a heuristic (interpolation) method *Int* across a wider y-axis range for comparison. Boxplots show the distribution of Mean Weighted Occupancy Errors (MOEs) for simulated tracks by analysis. Compared to Fig. S4, the MOE metric is the average error over all regions per track, weighted by the proportion of time spent in each region (14). For boxplot properties, see Fig. S4.

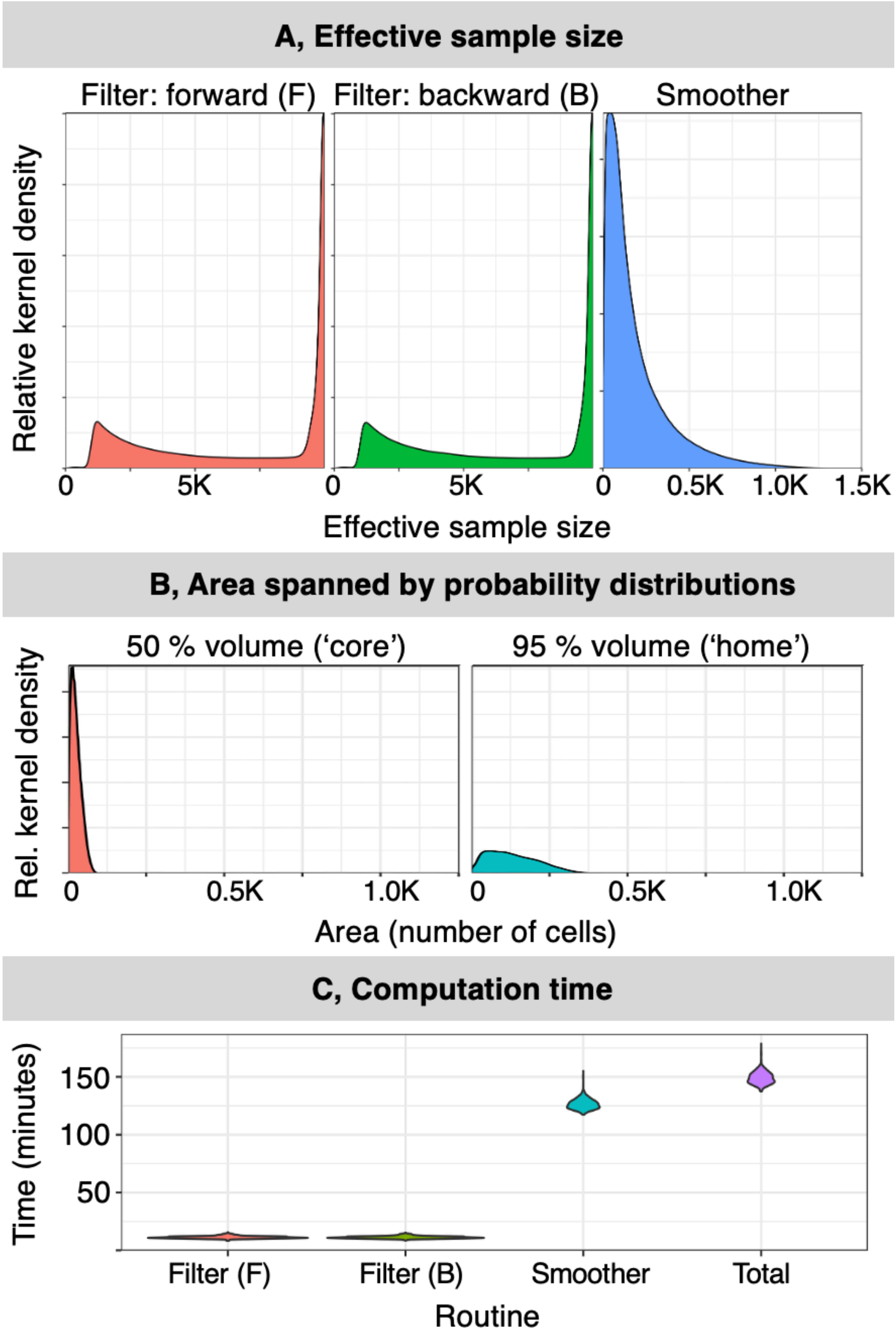


**Fig. S6. Simulation analysis: diagnostics. A** shows the distribution of effective sample sizes from the forward filter, backward filter and the two-filter smoother. **B** shows the distribution of areas spanned by smoothed probability distributions $f(\boldsymbol{s}_t \mid \boldsymbol{y}_{1:T})$. The smallest areas spanned by the 'core' of the distribution (encompassing 50 % of the probability mass) and the 'home range' (encompassing 95 % of the probability mass) are shown. **C** shows the distribution of computation times per routine/run. All statistics are shown for the 665/672 algorithm runs that converged (i.e., including 96 simulated individuals and both main and sensitivity analyses).

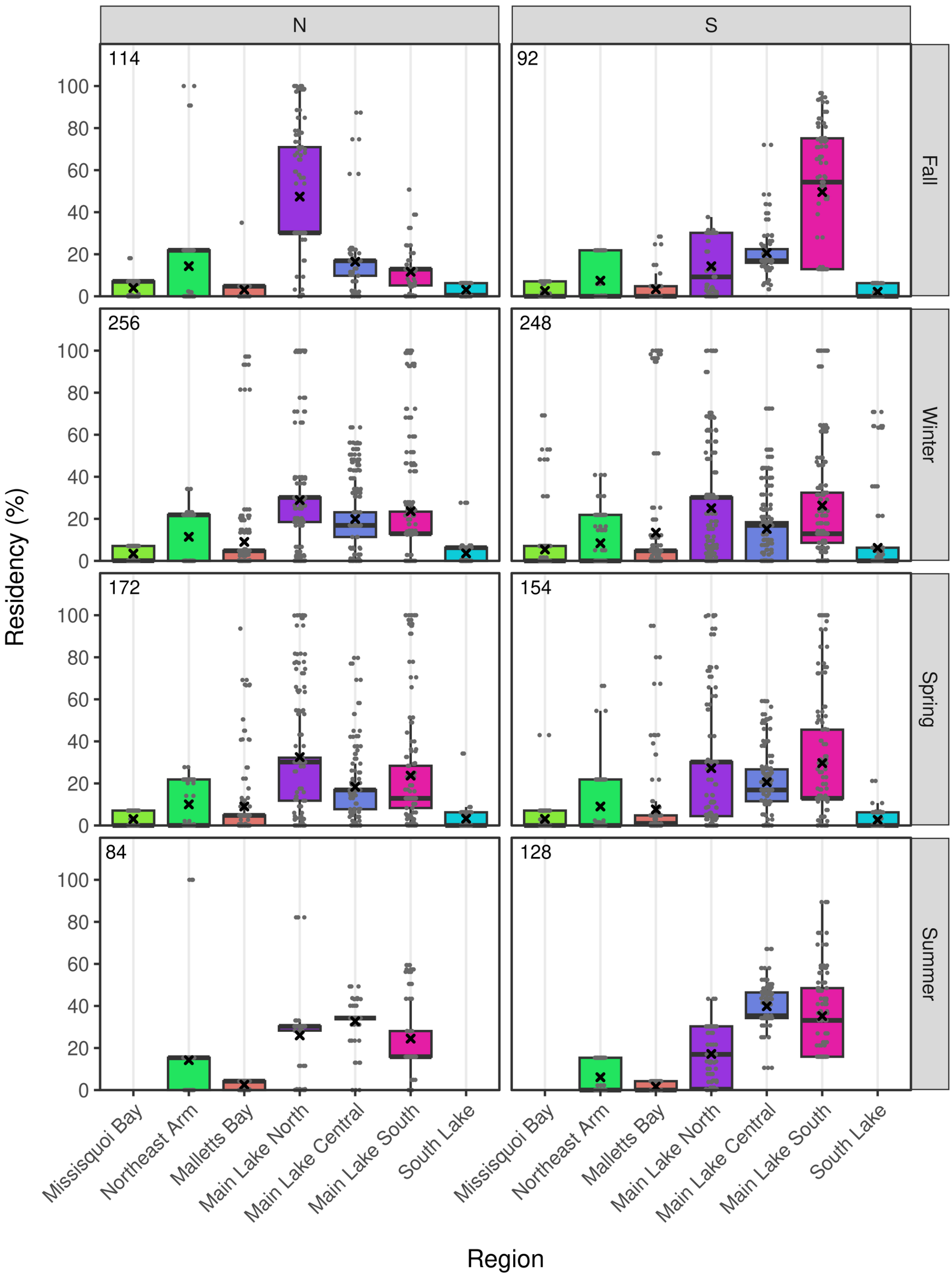


**Fig. S7. Real-world analysis: lake trout residency in Lake Champlain.** Panels show residency estimates for each region by tagging location (North, N or South, S) and season. Residency is the percentage of time spent in each region, derived by state-space modelling. (The real-world analysis did not include a heuristic comparison.) Small grey points mark jittered estimates. Boxplot quantiles are weighted by survival probability. Black crosses mark the weighted mean. The number of estimates is top left.

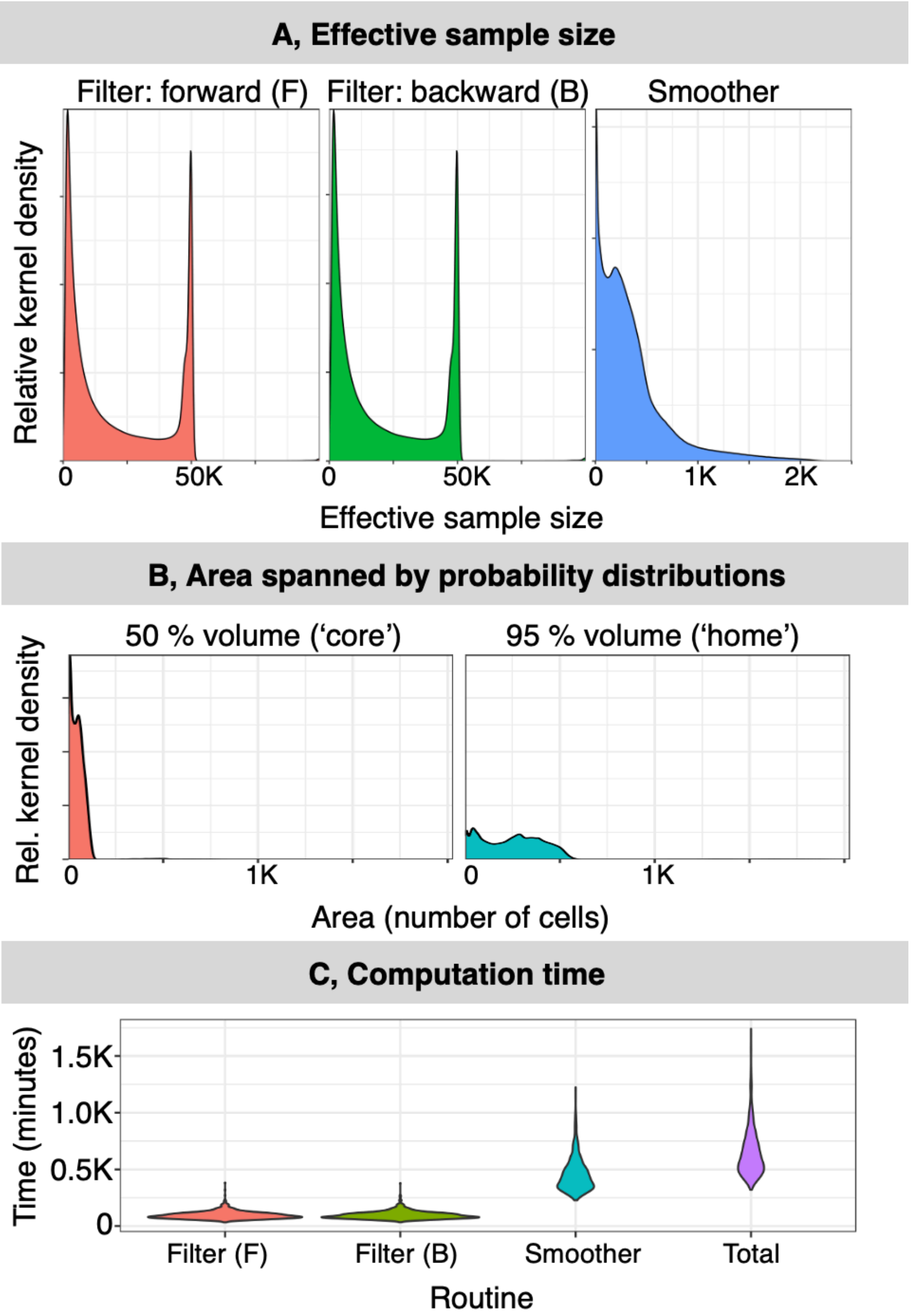


**Fig. S8. Real-world analysis: diagnostics, following Fig. S6.** For the real-world analysis, the acoustic time series were split into a set of 2070 overlapping blocks of approximately monthly duration (20,160–58,572 time steps). We performed locational inference using particle algorithms for the 1427 blocks with at least one detection in the surrounding blocks(s). (For the remaining blocks, we assumed a uniform distribution over the study area, accounting for summertime thermal habitat suitability.) This figure shows diagnostic statistics for the 1012/1427 blocks for which both the forward and backward particle filter runs and the two-filter smoother converged. Particle filters used 50,000–100,000 particles. The two-filter smoother used 2,500 particles.

**Tables**

**Table S1. A summary of tagged fish.** For each fish, the identifier, capture location and date, sex (male, M; female, F) and total length (mm) are shown.

| Row | ID | Longitude (°) | Latitude (°) | Date | Sex | Total length (mm) |
|---|---|---|---|---|---|---|
| 1 | 26784 | -73.3539 | 44.6881 | 2013-11-05 | M | 580 |
| 2 | 26788 | -73.3539 | 44.6881 | 2013-11-05 | F | 683 |
| 3 | 26789 | -73.3539 | 44.6881 | 2013-11-05 | F | 662 |
| 4 | 26790 | -73.3539 | 44.6881 | 2013-11-05 | F | 674 |
| 5 | 26791 | -73.3539 | 44.6881 | 2013-11-05 | M | 667 |
| 6 | 26792 | -73.3539 | 44.6881 | 2013-11-05 | F | 707 |
| 7 | 26793 | -73.3539 | 44.6881 | 2013-11-05 | F | 735 |
| 8 | 26794 | -73.3539 | 44.6881 | 2013-11-05 | M | 658 |
| 9 | 26795 | -73.3539 | 44.6881 | 2013-11-05 | F | 650 |
| 10 | 26796 | -73.3531 | 44.7015 | 2013-11-05 | M | 634 |
| 11 | 26797 | -73.3539 | 44.6881 | 2013-11-05 | F | 695 |
| 12 | 26800 | -73.3531 | 44.7015 | 2013-11-05 | M | 630 |
| 13 | 26801 | -73.3539 | 44.6881 | 2013-11-05 | M | 680 |
| 14 | 26802 | -73.3539 | 44.6881 | 2013-11-05 | F | 673 |
| 15 | 26803 | -73.3531 | 44.7015 | 2013-11-05 | M | 674 |
| 16 | 26804 | -73.3539 | 44.6881 | 2013-11-05 | F | 760 |
| 17 | 26805 | -73.3539 | 44.6881 | 2013-11-05 | M | 634 |
| 18 | 26806 | -73.3539 | 44.6881 | 2013-11-05 | F | 735 |
| 19 | 26807 | -73.3531 | 44.7015 | 2013-11-05 | M | 635 |
| 20 | 26808 | -73.3531 | 44.7015 | 2013-11-05 | M | 630 |
| 21 | 26770 | -73.3539 | 44.6881 | 2013-11-06 | M | 712 |
| 22 | 26771 | -73.3539 | 44.6881 | 2013-11-06 | F | 685 |
| 23 | 26773 | -73.3539 | 44.6881 | 2013-11-06 | F | 670 |
| 24 | 26774 | -73.3539 | 44.6881 | 2013-11-06 | M | 680 |
| 25 | 26778 | -73.3539 | 44.6881 | 2013-11-06 | M | 627 |
| 26 | 26785 | -73.3539 | 44.6881 | 2013-11-06 | F | 645 |
| 27 | 26786 | -73.3539 | 44.6881 | 2013-11-06 | F | 680 |
| 28 | 26787 | -73.3539 | 44.6881 | 2013-11-06 | M | 635 |
| 29 | 26799 | -73.3539 | 44.6881 | 2013-11-06 | M | 633 |
| 30 | 26776 | -73.3539 | 44.6881 | 2013-11-07 | F | 662 |
| 31 | 24320 | -73.3539 | 44.6876 | 2014-11-04 | M | 565 |
| 32 | 24321 | -73.3539 | 44.6876 | 2014-11-04 | M | 574 |
| 33 | 24322 | -73.3539 | 44.6876 | 2014-11-04 | M | 712 |
| 34 | 24323 | -73.3539 | 44.6876 | 2014-11-04 | M | 713 |
| 35 | 24324 | -73.3539 | 44.6876 | 2014-11-04 | M | 820 |
| 36 | 24325 | -73.3539 | 44.6876 | 2014-11-04 | M | 775 |
| 37 | 24326 | -73.3539 | 44.6876 | 2014-11-04 | F | 638 |
| 38 | 24327 | -73.3539 | 44.6876 | 2014-11-04 | M | 545 |
| 39 | 24328 | -73.3539 | 44.6876 | 2014-11-04 | M | 560 |
| 40 | 24329 | -73.3539 | 44.6876 | 2014-11-04 | F | 650 |
| 41 | 24330 | -73.3539 | 44.6876 | 2014-11-04 | M | 630 |
| 42 | 24331 | -73.3539 | 44.6876 | 2014-11-04 | F | 669 |

| Row | ID | Longitude (°) | Latitude (°) | Date | Sex | Total length (mm) |
|---|---|---|---|---|---|---|
| 43 | 24332 | -73.3539 | 44.6876 | 2014-11-04 | F | 677 |
| 44 | 24333 | -73.3539 | 44.6876 | 2014-11-04 | M | 651 |
| 45 | 24334 | -73.3539 | 44.6876 | 2014-11-04 | M | 558 |
| 46 | 24335 | -73.3539 | 44.6876 | 2014-11-05 | M | 632 |
| 47 | 24336 | -73.3530 | 44.7015 | 2014-11-05 | M | 672 |
| 48 | 24337 | -73.3539 | 44.6876 | 2014-11-05 | M | 680 |
| 49 | 24338 | -73.3539 | 44.6876 | 2014-11-05 | F | 679 |
| 50 | 24340 | -73.3539 | 44.6876 | 2014-11-05 | F | 705 |
| 51 | 24341 | -73.3539 | 44.6876 | 2014-11-05 | F | 709 |
| 52 | 24342 | -73.3539 | 44.6876 | 2014-11-05 | F | 720 |
| 53 | 24345 | -73.3539 | 44.6876 | 2014-11-05 | M | 612 |
| 54 | 24346 | -73.3539 | 44.6876 | 2014-11-05 | F | 626 |
| 55 | 24347 | -73.3539 | 44.6876 | 2014-11-05 | F | 637 |
| 56 | 24348 | -73.3539 | 44.6876 | 2014-11-05 | F | 658 |
| 57 | 24350 | -73.3539 | 44.6876 | 2014-11-05 | F | 616 |
| 58 | 24351 | -73.3539 | 44.6876 | 2014-11-05 | F | 670 |
| 59 | 24352 | -73.3539 | 44.6876 | 2014-11-05 | F | 680 |
| 60 | 24353 | -73.3539 | 44.6876 | 2014-11-05 | F | 649 |
| 61 | 24339 | -73.3315 | 44.2694 | 2014-11-10 | M | 712 |
| 62 | 24343 | -73.3315 | 44.2694 | 2014-11-10 | M | 665 |
| 63 | 24344 | -73.3315 | 44.2694 | 2014-11-10 | M | 674 |
| 64 | 24349 | -73.3315 | 44.2694 | 2014-11-10 | M | 683 |
| 65 | 24354 | -73.3315 | 44.2694 | 2014-11-10 | M | 685 |
| 66 | 24355 | -73.3315 | 44.2694 | 2014-11-10 | M | 685 |
| 67 | 24356 | -73.3315 | 44.2694 | 2014-11-10 | M | 705 |
| 68 | 24357 | -73.3315 | 44.2694 | 2014-11-10 | F | 720 |
| 69 | 24360 | -73.3078 | 44.2626 | 2014-11-10 | F | 648 |
| 70 | 24361 | -73.3315 | 44.2694 | 2014-11-10 | M | 630 |
| 71 | 24362 | -73.3315 | 44.2694 | 2014-11-10 | F | 695 |
| 72 | 24365 | -73.3078 | 44.2626 | 2014-11-10 | M | 645 |
| 73 | 24366 | -73.3315 | 44.2694 | 2014-11-10 | F | 675 |
| 74 | 24367 | -73.3315 | 44.2694 | 2014-11-10 | F | 750 |
| 75 | 24370 | -73.3315 | 44.2694 | 2014-11-10 | M | 617 |
| 76 | 24371 | -73.3315 | 44.2694 | 2014-11-10 | F | 627 |
| 77 | 24372 | -73.3315 | 44.2694 | 2014-11-10 | F | 714 |
| 78 | 24375 | -73.3315 | 44.2694 | 2014-11-10 | M | 678 |
| 79 | 24376 | -73.3315 | 44.2694 | 2014-11-10 | F | 665 |
| 80 | 24377 | -73.3315 | 44.2694 | 2014-11-10 | F | 675 |
| 81 | 24378 | -73.3315 | 44.2694 | 2014-11-10 | F | 635 |
| 82 | 24380 | -73.3078 | 44.2626 | 2014-11-10 | F | 710 |
| 83 | 24381 | -73.3315 | 44.2694 | 2014-11-10 | F | 709 |
| 84 | 24382 | -73.3315 | 44.2694 | 2014-11-10 | F | 710 |
| 85 | 24383 | -73.3315 | 44.2694 | 2014-11-10 | F | 683 |
| 86 | 24385 | -73.3078 | 44.2626 | 2014-11-10 | F | 655 |
| 87 | 24386 | -73.3315 | 44.2694 | 2014-11-10 | F | 752 |
| 88 | 24387 | -73.3315 | 44.2694 | 2014-11-10 | F | 715 |
| 89 | 24390 | -73.3315 | 44.2694 | 2014-11-10 | M | 680 |

| Row | ID | Longitude (°) | Latitude (°) | Date | Sex | Total length (mm) |
|---|---|---|---|---|---|---|
| 90 | 24391 | -73.3315 | 44.2694 | 2014-11-10 | M | 677 |
| 91 | 24392 | -73.3315 | 44.2694 | 2014-11-10 | M | 747 |
| 92 | 24393 | -73.3315 | 44.2694 | 2014-11-10 | F | 549 |
| 93 | 24394 | -73.3315 | 44.2694 | 2014-11-10 | M | 643 |

**Table S2. A summary of receiver deployments.** For each receiver deployment, an identifier is given alongside the station, receiver serial number, start and end dates, location and depth. Receiver locations are rounded for four decimal places. Rows sorted by station, start date and serial number.

| ID | Station | Receiver | Start | End | Longitude (°) | Latitude (°) | Depth (m) |
|---|---|---|---|---|---|---|---|
| 1 | Alburg | 125207 | 2014-08-21 | 2015-06-16 | -73.2740 | 44.8886 | 8.5 |
| 2 | Alburg | 125207 | 2015-06-16 | 2017-08-14 | -73.2737 | 44.8888 | 8.5 |
| 3 | Arnold Central | 125199 | 2014-08-11 | 2015-05-20 | -73.3890 | 44.1445 | 24.4 |
| 4 | Arnold Central | 125199 | 2015-05-20 | 2015-11-10 | -73.3889 | 44.1444 | 29.0 |
| 5 | Arnold Central | 125199 | 2015-11-10 | 2016-06-07 | -73.3888 | 44.1445 | 28.2 |
| 6 | Arnold Central | 125199 | 2016-06-07 | 2016-11-03 | -73.3886 | 44.1445 | 27.5 |
| 7 | Arnold Central | 125199 | 2016-11-03 | 2017-09-22 | -73.3888 | 44.1445 | 27.7 |
| 8 | Arnold East | 123545 | 2013-10-15 | 2013-11-20 | -73.3746 | 44.1509 | 15.5 |
| 9 | Arnold East | 123545 | 2013-11-20 | 2014-05-19 | -73.3749 | 44.1510 | 15.5 |
| 10 | Arnold East | 123545 | 2014-05-19 | 2014-08-11 | -73.3748 | 44.1510 | 15.5 |
| 11 | Arnold East | 123545 | 2014-08-11 | 2014-11-21 | -73.3750 | 44.1509 | 15.5 |
| 12 | Arnold East | 123545 | 2014-11-21 | 2015-05-20 | -73.3743 | 44.1506 | 15.5 |
| 13 | Arnold East | 123545 | 2015-05-20 | 2015-11-10 | -73.3742 | 44.1507 | 15.5 |
| 14 | Arnold East | 123545 | 2015-11-10 | 2016-06-07 | -73.3743 | 44.1506 | 15.1 |
| 15 | Arnold East | 123545 | 2016-06-07 | 2016-11-03 | -73.3746 | 44.1507 | 16.0 |
| 16 | Arnold East | 123545 | 2016-11-03 | 2017-07-27 | -73.3744 | 44.1506 | 15.0 |
| 17 | Arnold West | 125208 | 2014-08-11 | 2015-05-20 | -73.4091 | 44.1464 | 17.7 |
| 18 | Arnold West | 125208 | 2015-05-20 | 2015-11-10 | -73.4076 | 44.1463 | 21.0 |
| 19 | Arnold West | 125208 | 2015-11-10 | 2016-06-07 | -73.4076 | 44.1465 | 17.6 |
| 20 | Arnold West | 125208 | 2016-06-07 | 2016-11-03 | -73.4077 | 44.1464 | 17.6 |
| 21 | Arnold West | 125208 | 2016-11-03 | 2017-07-27 | -73.4077 | 44.1466 | 16.3 |
| 22 | Burlington | 123549 | 2013-10-08 | 2013-11-22 | -73.2323 | 44.4710 | 17.6 |
| 23 | Burlington | 123549 | 2013-11-22 | 2014-05-21 | -73.2322 | 44.4710 | 17.6 |
| 24 | Burlington | 123549 | 2014-05-21 | 2014-08-04 | -73.2320 | 44.4711 | 17.6 |
| 25 | Burlington | 123549 | 2014-08-21 | 2014-11-23 | -73.2319 | 44.4710 | 17.6 |
| 26 | Burlington | 123549 | 2014-11-23 | 2015-05-22 | -73.2321 | 44.4711 | 17.6 |
| 27 | Burlington | 123549 | 2015-05-22 | 2015-11-11 | -73.2321 | 44.4711 | 30.0 |

| ID | Station | Receiver | Start | End | Longitude (°) | Latitude (°) | Depth (m) |
|---|---|---|---|---|---|---|---|
| 28 | Burlington | 123549 | 2015-11-11 | 2016-06-24 | -73.2321 | 44.4711 | 18.0 |
| 29 | Burlington | 123549 | 2016-06-24 | 2016-10-26 | -73.2321 | 44.4711 | 18.0 |
| 30 | Burlington | 123549 | 2016-10-26 | 2017-08-10 | -73.2324 | 44.4712 | 18.2 |
| 31 | Carry | 125210 | 2014-08-21 | 2015-06-16 | -73.2910 | 44.8452 | 6.7 |
| 32 | Carry | 125210 | 2015-06-16 | 2016-05-19 | -73.2912 | 44.8451 | 6.7 |
| 33 | Carry | 125210 | 2016-05-19 | 2016-11-02 | -73.2912 | 44.8449 | 6.0 |
| 34 | Carry | 125210 | 2016-11-02 | 2017-07-12 | -73.2912 | 44.8449 | 5.7 |
| 35 | Crown Point | 125213 | 2014-08-11 | 2015-05-20 | -73.4099 | 44.0231 | 7.6 |
| 36 | Crown Point | 125213 | 2015-05-20 | 2016-06-07 | -73.4100 | 44.0232 | 7.0 |
| 37 | Gordon | 123540 | 2013-10-17 | 2013-11-22 | -73.3539 | 44.6881 | 20.0 |
| 38 | Gordon | 123540 | 2013-11-22 | 2014-05-20 | -73.3539 | 44.6881 | 6.3 |
| 39 | Gordon | 123540 | 2014-05-20 | 2014-11-23 | -73.3539 | 44.6876 | 6.3 |
| 40 | Gordon | 123540 | 2014-11-23 | 2015-06-17 | -73.3540 | 44.6877 | 6.3 |
| 41 | Gordon | 123540 | 2015-06-17 | 2015-11-17 | -73.3539 | 44.6874 | 21.0 |
| 42 | Gordon | 123540 | 2015-11-17 | 2016-06-17 | -73.3539 | 44.6874 | 21.0 |
| 43 | Gordon | 123540 | 2016-06-17 | 2016-11-02 | -73.3539 | 44.6874 | 21.0 |
| 44 | Gordon | 123540 | 2016-11-02 | 2017-08-14 | -73.3539 | 44.6875 | 20.8 |
| 45 | Gut | 123538 | 2013-10-17 | 2013-11-22 | -73.3014 | 44.7595 | 3.4 |
| 46 | Gut | 123538 | 2013-11-22 | 2014-05-20 | -73.3015 | 44.7593 | 6.3 |
| 47 | Gut | 123538 | 2014-05-20 | 2015-05-21 | -73.3021 | 44.7591 | 4.5 |
| 48 | Gut | 123538 | 2015-05-21 | 2016-05-19 | -73.3024 | 44.7591 | 4.5 |
| 49 | Gut | 123538 | 2016-05-19 | 2017-06-23 | -73.3024 | 44.7592 | 4.5 |
| 50 | Hog Island | 125204 | 2014-08-21 | 2015-06-16 | -73.2285 | 44.9242 | 6.1 |
| 51 | Hog Island | 125204 | 2015-06-16 | 2016-05-19 | -73.2283 | 44.9242 | 6.1 |
| 52 | Hog Island | 125204 | 2016-05-19 | 2016-11-02 | -73.2280 | 44.9246 | 7.0 |
| 53 | Hog Island | 125204 | 2016-11-02 | 2017-07-12 | -73.2283 | 44.9245 | 6.7 |
| 54 | Island Line East | 123546 | 2013-10-15 | 2013-11-22 | -73.3051 | 44.5894 | 5.2 |
| 55 | Island Line East | 123546 | 2013-11-22 | 2014-05-21 | -73.3051 | 44.5896 | 5.0 |
| 56 | Island Line East | 123546 | 2014-05-21 | 2015-06-17 | -73.3049 | 44.5898 | 5.0 |
| 57 | Island Line East | 123546 | 2015-06-17 | 2016-06-24 | -73.3049 | 44.5898 | 5.0 |

| ID | Station | Receiver | Start | End | Longitude (°) | Latitude (°) | Depth (m) |
|---|---|---|---|---|---|---|---|
| 58 | Island Line East | 123546 | 2016-06-24 | 2016-11-08 | -73.3048 | 44.5899 | 5.0 |
| 59 | Island Line East | 123547 | 2016-11-08 | 2017-08-01 | -73.3048 | 44.5899 | 5.0 |
| 60 | Island Line West | 123539 | 2013-10-15 | 2013-11-22 | -73.3231 | 44.5966 | 5.0 |
| 61 | Island Line West | 123539 | 2013-11-22 | 2014-05-21 | -73.3233 | 44.5967 | 5.2 |
| 62 | Island Line West | 123539 | 2014-05-21 | 2015-06-17 | -73.3233 | 44.5966 | 5.2 |
| 63 | Island Line West | 123539 | 2015-06-17 | 2016-06-24 | -73.3232 | 44.5966 | 5.2 |
| 64 | Island Line West | 123539 | 2016-06-24 | 2016-11-02 | -73.3232 | 44.5966 | 5.8 |
| 65 | Island Line West | 123539 | 2016-11-02 | 2017-08-01 | -73.3232 | 44.5966 | 5.6 |
| 66 | Isle LaMotte | 125203 | 2014-08-21 | 2015-06-16 | -73.3114 | 44.8898 | 5.2 |
| 67 | Isle LaMotte | 125203 | 2015-06-16 | 2016-05-19 | -73.3114 | 44.8893 | 6.0 |
| 68 | Isle LaMotte | 125203 | 2016-05-19 | 2016-11-02 | -73.3115 | 44.8891 | 7.0 |
| 69 | Isle LaMotte | 125203 | 2016-11-02 | 2017-07-12 | -73.3118 | 44.8889 | 6.7 |
| 70 | Malletts Bay | 130644 | 2016-11-18 | 2017-07-26 | -73.2445 | 44.6078 | - |
| 71 | Missisquoi | 123548 | 2013-10-30 | 2013-11-22 | -73.2239 | 44.9653 | 4.8 |
| 72 | Missisquoi | 123548 | 2013-11-22 | 2014-05-22 | -73.2240 | 44.9651 | 4.8 |
| 73 | Point Au Roche | 125202 | 2014-08-21 | 2015-06-16 | -73.3648 | 44.8272 | 21.3 |
| 74 | Point Au Roche | 125202 | 2015-06-16 | 2016-05-19 | -73.3651 | 44.8271 | 25.0 |
| 75 | Point Au Roche | 125202 | 2016-05-19 | 2016-11-02 | -73.3650 | 44.8277 | 24.0 |
| 76 | Point Au Roche | 125202 | 2016-11-02 | 2017-07-12 | -73.3650 | 44.8275 | 23.7 |
| 77 | Rockwell | 123541 | 2013-10-17 | 2013-11-22 | -73.3539 | 44.6676 | 5.0 |
| 78 | Rockwell | 123541 | 2013-11-22 | 2014-05-20 | -73.3539 | 44.6676 | 20.0 |
| 79 | Rockwell | 123541 | 2014-05-20 | 2014-11-23 | -73.3538 | 44.6673 | 5.0 |
| 80 | Rockwell | 123541 | 2014-11-23 | 2015-06-17 | -73.3539 | 44.6674 | 5.0 |
| 81 | Rockwell | 123541 | 2015-06-17 | 2015-11-17 | -73.3539 | 44.6674 | 5.0 |
| 82 | Rockwell | 123541 | 2015-11-17 | 2016-06-17 | -73.3539 | 44.6674 | 5.0 |
| 83 | Rockwell | 123541 | 2016-06-17 | 2016-11-02 | -73.3539 | 44.6674 | 6.1 |
| 84 | Rockwell | 123541 | 2016-11-02 | 2017-07-26 | -73.3541 | 44.6674 | 7.0 |
| 85 | Rouses Point | 125214 | 2014-08-21 | 2015-06-16 | -73.3472 | 45.0029 | 5.2 |
| 86 | Rouses Point | 125214 | 2015-06-16 | 2016-05-19 | -73.3473 | 45.0028 | 9.0 |
| 87 | Rouses Point | 125214 | 2016-05-19 | 2016-11-02 | -73.3474 | 45.0028 | 9.0 |

| ID | Station | Receiver | Start | End | Longitude (°) | Latitude (°) | Depth (m) |
|---|---|---|---|---|---|---|---|
| 88 | Rouses Point | 125214 | 2016-11-02 | 2017-07-12 | -73.3475 | 45.0028 | 7.2 |
| 89 | Sand Bar Inland Sea | 123544 | 2013-10-17 | 2013-11-22 | -73.2500 | 44.6356 | 6.3 |
| 90 | Sand Bar Inland Sea | 123544 | 2013-11-22 | 2014-05-20 | -73.2506 | 44.6356 | 9.0 |
| 91 | Sand Bar Inland Sea | 123544 | 2014-05-20 | 2015-05-21 | -73.2507 | 44.6358 | 9.0 |
| 92 | Sand Bar Inland Sea | 123544 | 2015-05-21 | 2016-05-19 | -73.2505 | 44.6358 | 9.0 |
| 93 | Sand Bar Inland Sea | 123544 | 2016-05-19 | 2016-11-08 | -73.2505 | 44.6359 | 9.0 |
| 94 | Sand Bar Inland Sea | 125213 | 2016-11-08 | 2017-07-07 | -73.2505 | 44.6359 | 9.0 |
| 95 | Sand Bar Malletts | 123543 | 2013-10-15 | 2014-05-21 | -73.2616 | 44.6174 | 4.8 |
| 96 | Sand Bar Malletts | 123543 | 2014-05-21 | 2015-06-17 | -73.2614 | 44.6175 | 4.8 |
| 97 | Sand Bar Malletts | 123543 | 2015-06-17 | 2016-06-24 | -73.2614 | 44.6175 | 4.8 |
| 98 | Sand Bar Malletts | 123543 | 2016-06-24 | 2016-11-08 | -73.2614 | 44.6175 | 4.8 |
| 99 | Sand Bar Malletts | 125205 | 2016-11-08 | 2017-08-01 | -73.2614 | 44.6175 | 4.8 |
| 100 | Saxton | 125201 | 2014-08-11 | 2014-11-21 | -73.2825 | 44.3929 | 16.2 |
| 101 | Saxton | 125201 | 2014-11-21 | 2015-05-22 | -73.2825 | 44.3925 | 17.1 |
| 102 | Saxton | 125201 | 2015-05-22 | 2015-11-10 | -73.2825 | 44.3925 | 17.1 |
| 103 | Saxton | 125201 | 2015-11-10 | 2016-06-07 | -73.2825 | 44.3926 | 17.1 |
| 104 | Saxton | 125201 | 2016-06-07 | 2016-11-03 | -73.2823 | 44.3928 | 17.7 |
| 105 | Saxton | 125201 | 2016-11-03 | 2017-07-20 | -73.2823 | 44.3928 | 16.7 |
| 106 | Schuyler | 125200 | 2014-08-21 | 2014-11-21 | -73.3774 | 44.5124 | 18.3 |
| 107 | Schuyler | 125200 | 2014-11-21 | 2015-05-22 | -73.3774 | 44.5127 | 18.1 |
| 108 | Schuyler | 125200 | 2015-05-22 | 2015-11-11 | -73.3776 | 44.5128 | 18.1 |
| 109 | Schuyler | 125200 | 2015-11-11 | 2016-06-10 | -73.3777 | 44.5127 | 18.1 |
| 110 | Schuyler | 125200 | 2016-06-10 | 2016-10-26 | -73.3778 | 44.5127 | 18.5 |
| 111 | Schuyler | 125200 | 2016-10-26 | 2017-07-20 | -73.3777 | 44.5128 | 18.0 |
| 112 | Shelburne | 125206 | 2014-08-11 | 2014-11-21 | -73.2464 | 44.4441 | 9.1 |
| 113 | Shelburne | 125206 | 2014-11-21 | 2015-05-22 | -73.2465 | 44.4441 | 30.0 |
| 114 | Shelburne | 125206 | 2015-05-22 | 2015-08-04 | -73.2464 | 44.4441 | 12.7 |
| 115 | Shelburne | 125206 | 2015-08-04 | 2015-11-11 | -73.2464 | 44.4441 | 12.7 |
| 116 | Shelburne | 125206 | 2015-11-11 | 2016-06-10 | -73.2464 | 44.4441 | 12.7 |
| 117 | Shelburne | 125206 | 2016-06-10 | 2016-10-26 | -73.2465 | 44.4442 | 12.8 |

| ID | Station | Receiver | Start | End | Longitude (°) | Latitude (°) | Depth (m) |
|---|---|---|---|---|---|---|---|
| 118 | Shelburne | 125206 | 2016-10-26 | 2017-07-20 | -73.2463 | 44.4444 | 12.9 |
| 119 | Split Rock | 125205 | 2014-08-11 | 2015-05-20 | -73.3078 | 44.2626 | 17.7 |
| 120 | Split Rock | 125205 | 2015-05-20 | 2015-11-25 | -73.3082 | 44.2623 | 18.0 |
| 121 | Split Rock | 125211 | 2015-11-25 | 2017-09-22 | -73.3082 | 44.2623 | 18.0 |
| 122 | Trembleau | 125212 | 2014-08-21 | 2015-06-16 | -73.3678 | 44.8785 | 10.1 |
| 123 | Trembleau | 125212 | 2015-06-16 | 2016-05-19 | -73.3676 | 44.8784 | 11.0 |
| 124 | Trembleau | 125212 | 2016-05-19 | 2016-11-02 | -73.3677 | 44.8786 | 11.0 |
| 125 | Trembleau | 125212 | 2016-11-02 | 2017-07-12 | -73.3678 | 44.8786 | 10.7 |
| 126 | Whallon | 123542 | 2013-10-15 | 2013-11-20 | -73.3316 | 44.2693 | 17.1 |
| 127 | Whallon | 123542 | 2013-11-20 | 2014-05-19 | -73.3319 | 44.2694 | 17.1 |
| 128 | Whallon | 123542 | 2014-05-19 | 2014-11-21 | -73.3315 | 44.2694 | 19.0 |
| 129 | Whallon | 123542 | 2014-11-21 | 2015-05-20 | -73.3312 | 44.2695 | 19.0 |
| 130 | Whallon | 123542 | 2015-05-20 | 2015-11-10 | -73.3311 | 44.2694 | 19.0 |
| 131 | Whallon | 123542 | 2015-11-10 | 2016-06-07 | -73.3310 | 44.2695 | 19.1 |
| 132 | Whallon | 123542 | 2016-06-07 | 2016-11-03 | -73.3311 | 44.2694 | 19.1 |
| 133 | Whallon | 123542 | 2016-11-03 | 2017-07-27 | -73.3311 | 44.2694 | 19.0 |
| 134 | Wilcox | 123547 | 2013-10-23 | 2013-11-22 | -73.3529 | 44.7016 | 6.3 |
| 135 | Wilcox | 123547 | 2013-11-22 | 2014-05-20 | -73.3531 | 44.7015 | 3.4 |
| 136 | Wilcox | 123547 | 2014-05-20 | 2014-09-25 | -73.3530 | 44.7015 | 6.3 |
| 137 | Wilcox | 123547 | 2014-09-25 | 2014-11-23 | -73.3530 | 44.7015 | 6.3 |
| 138 | Wilcox | 123547 | 2014-11-23 | 2015-05-21 | -73.3530 | 44.7015 | 6.3 |
| 139 | Wilcox | 123547 | 2015-05-21 | 2015-11-17 | -73.3529 | 44.7016 | 6.3 |
| 140 | Wilcox | 123547 | 2015-11-17 | 2016-06-17 | -73.3529 | 44.7016 | 6.3 |
| 141 | Wilcox | 123547 | 2016-06-17 | 2016-08-10 | -73.3528 | 44.7016 | 6.3 |
| 142 | Willsboro | 125209 | 2014-08-21 | 2014-11-21 | -73.3930 | 44.4403 | 45.7 |
| 143 | Willsboro | 125209 | 2014-11-21 | 2015-05-22 | -73.3927 | 44.4404 | 45.7 |
| 144 | Willsboro | 125209 | 2015-05-22 | 2015-11-11 | -73.3925 | 44.4408 | 45.7 |
| 145 | Willsboro | 125209 | 2015-11-11 | 2016-06-10 | -73.3926 | 44.4404 | 39.1 |
| 146 | Willsboro | 125209 | 2016-06-10 | 2016-10-26 | -73.3924 | 44.4402 | 38.6 |
| 147 | Willsboro | 125209 | 2016-10-26 | 2017-07-20 | -73.3923 | 44.4402 | 37.3 |

| ID | Station | Receiver | Start | End | Longitude (°) | Latitude (°) | Depth (m) |
|---|---|---|---|---|---|---|---|
| 148 | Winooski Delta Mid | 127130 | 2015-09-21 | 2016-07-11 | -73.2898 | 44.5213 | - |
| 149 | Winooski Delta Mid | 127130 | 2016-09-01 | 2017-08-29 | -73.2898 | 44.5213 | - |
| 150 | Winooski Delta North | 127337 | 2015-09-21 | 2016-07-01 | -73.3132 | 44.5513 | - |
| 151 | Winooski Delta North | 127337 | 2016-09-01 | 2017-08-29 | -73.3132 | 44.5513 | - |
| 152 | Winooski Delta South | 127129 | 2015-09-21 | 2016-07-11 | -73.2803 | 44.5070 | - |
| 153 | Winooski Delta South | 127129 | 2016-09-01 | 2017-08-29 | -73.2755 | 44.4952 | - |

**Table S3. A summary of ancillary datasets.** We sourced and analysed ancillary datasets to inform the design of our movement and acoustic observation models. For each model, the names of the datasets we analysed, alongside information on location, source and the model components informed by our analysis, is listed. Model formulation was also informed by wider consideration of the literature and domain expertise.

| Model | Dataset | Location | Source | Components |
|---|---|---|---|---|
| Movement model | Alexie Accelerometery | Alexie Lake, Northwest Territories, Canada | (24, 25) | Step lengths |
| | Drummond Island VPS | Lake Huron, Michigan, USA | (26) | Step lengths, turning angles |
| | Thunder Bay VPS | Lake Huron, Michigan, USA | (29) | Step lengths, turning angles |
| | Champlain Range Test | Lake Champlain | (7) | Detection probability |
| | Ontario Range Test | St Lawrence Channel, Lake Ontario, USA/Canada | (30) | Detection probability |

**Table S4. A summary of parameter values.** We conducted simulation and real-world analyses using our best model. In the simulation sensitivity analysis, we also explored the effects of restrictive (-) and flexible (+) model parameterisations. In each adjusted parameterisation, we adjusted one of the model components while holding other components constant at the values used in the best model.

| **Row** | **Category** | **Movement model** | | | | **Acoustic observation model** | | |
|---|---|---|---|---|---|---|---|---|
| | | **Step length, $d$** | | | **Turning angle, $\Delta\phi$** | | | |
| | | **mobility** | $\boldsymbol{k}$ | $\boldsymbol{\theta}$ | $\boldsymbol{\sigma}$ | $\boldsymbol{\alpha}$ | $\boldsymbol{\beta}$ | $\boldsymbol{\gamma}$ |
| 1 | Best | 216 | 3.2500 | 25.0000 | 0.4000 | 0.9039 | -0.0021 | 7000 |
| 2 | Step(-) | 162 | 4.3333 | 16.8750 | 0.4000 | 0.9039 | -0.0021 | 7000 |
| 3 | Step(+) | 270 | 2.6000 | 35.1562 | 0.4000 | 0.9039 | -0.0021 | 7000 |
| 4 | Angle(-) | 216 | 3.2500 | 25.0000 | 0.3000 | 0.9039 | -0.0021 | 7000 |
| 5 | Angle(+) | 216 | 3.2500 | 25.0000 | 0.5000 | 0.9039 | -0.0021 | 7000 |
| 6 | AC(-) | 216 | 3.2500 | 25.0000 | 0.4000 | 0.6779 | -0.0026 | 5250 |
| 7 | AC(+) | 216 | 3.2500 | 25.0000 | 0.4000 | 1.1299 | -0.0016 | 8750 |

1141
